# Molecular switch architecture drives response properties

**DOI:** 10.1101/2020.06.12.147900

**Authors:** Khem Raj Ghusinga, Roger D. Jones, Alan M. Jones, Timothy C. Elston

## Abstract

Many intracellular signaling pathways are composed of molecular switches, proteins that transition between two states—*on* and *off*. Typically, signaling is initiated when an external stimulus activates its cognate receptor that in turn causes downstream switches to transition from *off* to *on* using one of the following mechanisms: activation, in which the transition rate from the *off* state to the *on* state increases; derepression, in which the transition rate from the *on* state to the *off* state decreases; and concerted, in which activation and derepression operate simultaneously. We use mathematical modeling to compare these signaling mechanisms in terms of their dose-response curves, response times, and abilities to process upstream fluctuations. Our analysis elucidates several general principles. First, activation increases the sensitivity of the pathway, whereas derepression decreases sensitivity. Second, activation generates response times that decrease with signal strength, whereas derepression causes response times to increase with signal strength. These opposing features allow the concerted mechanism to not only show dose-response alignment, but also to decouple the response time from stimulus strength. However, these potentially beneficial properties come at the expense of increased susceptibility to upstream fluctuations. In addition to above response metrics, we also examine the effect of receptor removal on switches governed by activation and derepression. We find that if inactive (active) receptors are preferentially removed then activation (derepression) exhibits a sustained response whereas derepression (activation) adapts. In total, we show how the architecture of molecular switches govern their response properties. We also discuss the biological implications of our findings.

## Introduction

Several molecules involved in intracellular signaling pathways act as molecular switches. These are proteins that can be temporarily modified to transition between two conformations, one corresponding to an *on* (active) state and another to an *off* (inactive) state. Two prominent examples of such switches are proteins that are modified by phosphorylation and dephosphorylation and GTPases that bind nucleotides. For phosphorylation-dephosphorylation cycles, it is common for the covalent addition of a phosphate by a kinase to cause activation of the modified protein. A phosphatase removes the phosphate to turn the protein *off*. In the GTPase cycle, the protein is *on* when bound to Guanosine triphosphate (GTP) and *off* when bound to Guanosine diphosphate (GDP). The transition from the GDP bound state to the GTP bound state requires nucleotide exchange, whereas the transition from GTP bound to GDP bound state is achieved via hydrolysis of the *γ* phosphate on GTP. The basal rates of nucleotide exchange and hydrolysis are often small. These reaction rates are increased several fold by Guanine Exchange Factors (GEFs) and GTPase Accelerating Proteins (GAPs), respectively [1, 2].

A signaling pathway is often initiated upon recognition of a stimulus by its cognate receptor, which then activates a downstream switch. In principle, a switch may be turned *on* by *at least* two mechanisms: a) by increasing the transition rate from the *off* state to the *on* state, and b) by decreasing the transition rate from the *on* state to the *off* state. We term these mechanisms *activation* and *derepression*, respectively. Examples of both these mechanisms are found in the GTPase cycle. In animals, signaling through many pathways is initiated by G protein coupled receptors (GPCRs) that respond to a diverse set of external stimuli. These receptors act as GEFs to activate heterotrimeric G proteins [3–6]. Thus, pathway *activation* relies upon increasing the transition rate from the *off* state to the *on* state. There are no GPCRs in plants and other bikonts; the nucleotide exchange occurs spontaneously, without requiring GEF activity [7–9]. G proteins are kept in the *off* state by a repressor such as a GAP or some other protein that holds the self-activating G protein in its inactive state. In this scenario, the presence of a stimulus results in *derepression*, i.e., removal of the repressing activity [10–12]. These two mechanisms for initiating signaling, activation and derepression, are not mutually exclusive. For example, a *concerted* signal initiation, whereby both activation and dererpression are used, is employed in the GTPase cycle of the yeast mating response pathway [13,14]. In this example, inactive GPCRs recruit a GAP protein and act to repress, whereas active receptors have GEF activity and act to activate. Thus, perception of a stimulus leads to concerted activation and derepression by increasing GEF activity while decreasing GAP activity.

These three mechanisms are not limited to GTPase cycles. The activation mechanism described here in fact is a simpler abstraction of a linear signaling cascade, a classical framework used to study general properties of signaling pathways [15–19] as well as to model specific signaling pathways [20–22]. While derepression may seem like an unusual mechanism, it occurs in numerous important signaling pathways in plants (e.g., auxin, ethylene, gibberellin, phytochrome), as well as gene regulation [23–27]. In many of these cases, derepression occurs through decrease in the degradation rate of a component instead of its deactivation rate. Concerted mechanisms are found in bacterial two component systems, wherein the same component acts as kinase and phosphatase [28–35].

Mathematical modeling has proven to be a useful tool for understanding the design principles of signaling pathways, and, not surprisingly, mathematical models of activation, derepression and concerted mechanisms have been studied previously. For example, the classical Goldbeter-Koshland model studied zero-order ultrasensitivity of an activation mechanism [15]. Further analyses have examined the effect of receptor numbers [36–38], feedback mechanisms [39, 40], removal of active receptors via endocytosis and degradation [41, 42], etc. Similarly, important properties of the concerted mechanism, such as its ability to do ratiometric signaling [13, 14], to align dose responses at different stages of the signaling pathway [43], as well as its robustness [29, 44] are well-known. The derepression model is relatively less studied. Although there are models of G-signaling in *Arabidopsis thaliana* [45–47], these models have a large number of states and parameters and do not specifically look at properties of derepression mechanism.

Despite these efforts, a systematic comparison of various properties of activation, derepression, and concerted mechanisms of signaling has been lacking. Comparing these mechanisms should enable our understanding of why different organisms have chosen different mechanisms. To this end, we specifically choose four metrics for the comparison: a) dose-response, b) response time, c) ability to suppress or filter stochastic fluctuations in upstream components, and d) effect of receptor removal. The rationale behind comparing dose response curves is that they provide information about the input sensitivity range and the output dynamic range, both of which are of pharmacological importance. We supplement this comparison with response times, which provide information about the dynamics of the signaling activity. The third metric of comparison is motivated from the fact that signaling pathways are subject to inherent stochastic nature of biochemical reactions, further compounded by fluctuations in the number of components [48–53]. Lastly, we study the effect of receptor removal on the response of these signaling mechanisms because many signaling pathways evince receptor removal [11, 42, 54–56]. We study these properties by constructing both deterministic ordinary differential equation models and stochastic models based on continuous-time Markov chains.

Our results show that activation has the following two effects: it makes the switch response more sensitive than that of the receptor, and it speeds up the response with the stimulus strength. In contrast, derepression makes the switch response less sensitive than the receptor occupancy and slows down the response speed as stimulus strength increases. These counteracting behaviors of activation and derepression lead to intermediate sensitivity and intermediate response time for the concerted mechanism. In the special case of a perfect concerted mechanism (equal activation and repression), the dose-response curve of the pathway aligns with the receptor occupancy and the response time does not depend upon the stimulus level. The noise comparison reveals that the concerted mechanism is more susceptible to fluctuations than the activation and derepression mechanisms, which perform similarly. Finally, our analysis of the effect of receptor removal highlights another important difference between activation and derepression. Removal of active (inactive) receptors at a faster rate than inactive (active) receptors results in an adaptive response for activation (derepression) and sustained response for derepression (activation). We finally compare our findings with experimental observations, suggesting reasons that might have led biological systems to choose one of these mechanisms over the others.

## Model formulation

We consider a two-tier model for each of three mechanisms of signaling through a molecular switch (Fig. 1). The first tier is common for all mechanisms, where an inactive receptor (*X*) becomes active (*X*\*) when its corresponding input (stimulus) is presented. The second tier is the molecular switch that transitions between *off* (*Y*) and *on* (*X*\*) states. In the activation mechanism, the transition rate from the *off* state to the *on* state increases as the number of active receptor molecules increases (Fig. 1(a)). In the derepression mechanism, the transition rate from the *on* state to the *off* decreases with decrease in the number of inactive receptor molecules (Fig. 1(b)). In the concerted mechanism, both activation and derepression occur simultaneously (Fig. 1(c)). We model these mechanisms using ordinary differential equations (ODEs), assuming mass-action kinetics. To this end, we denote the time by *t*, stimulus level by *S*, the total number of receptors by *X_T_*, and the total number of switches by *Y_T_*. We use *X*\* and *Y*\* to denote the number of active receptors and the number of active switches, respectively. The rate constants are as follows: *k*_1_ is the rate of receptor activation per unit stimulus, *k*_2_ is the rate of receptor deactivation, *k*_3_ is the basal rate of activation of the switch, *k*_4_ is the basal rate of deactivation of the switch, *k*_5_ is the strength of activation of an individual active receptor, and *k*_6_ is the strength of repression of an individual inactive receptor. Thus, the (total) activation strength is *k*_5_*X_T_* and (total) repression strength is *k*_6_*X_T_*. Lastly, we assume that *X_T_* and *Y_T_* are conserved, and that each model is in steady state before presentation of the stimulus at *t* = 0.

**Figure 1:**
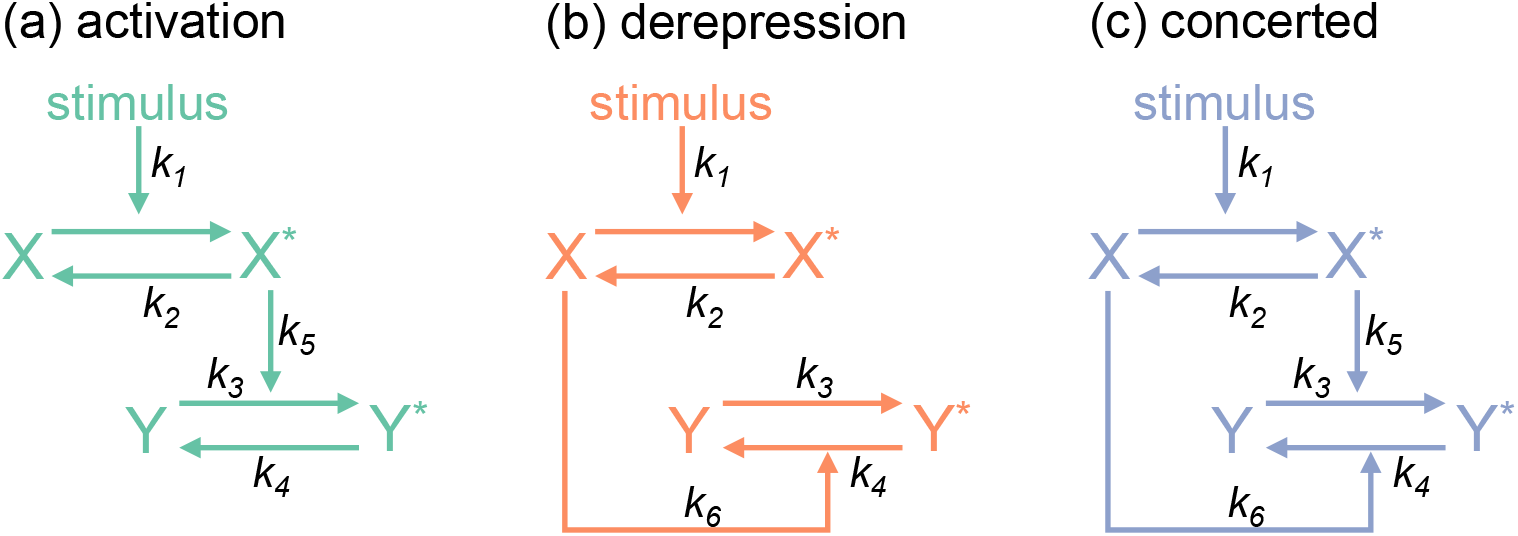
Mechanisms for signaling through molecular switches. Presentation of a stimulus activates a receptor (*X* → *X*\*). The reverse reaction causes deactivation of the receptor (*X*\* → *X*). These transitions govern the activity of a molecular switch downstream. (**a**) In the activation mechanism, *X*\* increases the rate at which the inactive switch (*Y*) becomes active (*Y*\*). The opposite reaction *Y*\* → *Y* has a constant rate. (**b**) In the derepression mechanism, the transition *Y* → *Y*\* occurs at a constant rate. Activity of the switch is controlled through *X*: the stimulus decreases *X* and consequently increases *Y*\*. (**c**) In the concerted paradigm, both activation and derepression simultaneously control the downstream component.

Note that the concerted mechanism encompasses both activation and derepression. Therefore, writing ODEs for the concerted mechanism is sufficient to capture all three mechanisms. The number of active receptors and the number of active switches evolve over time according to the ODEs:

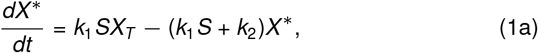

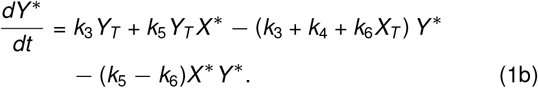

The activation and derepression mechanisms represent limiting cases in which *k*_6_ = 0 and *k*_5_ = 0, respectively. Solving (1) requires rate constants and initial conditions to be specified. We assume that initial conditions are given by the pre-stimulus (*S* = 0) steady state:

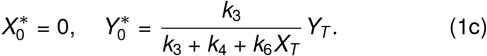

With the models described by (1), we next compare the three signaling mechanisms in terms of their dose responses and response times.

## Dose responses

We begin our analysis by examining the steady-state dose responses of activation, derepression, and concerted mechanisms. The steady-state solution to (1) is

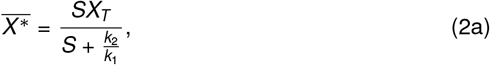

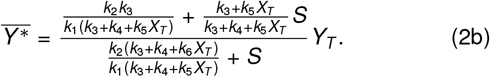

Here 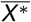 and 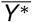 are the number of active (occupied) receptors and the number of active switches long time after the stimulus is presented (*t* → ∞), respectively. Notably, both 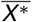 and 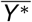 have the form

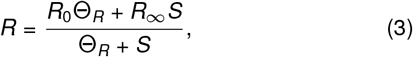

where *R*_0_ is the minimum response corresponding to *S* = 0, *R_∞_* is the maximum response corresponding to *S* ≫ Θ_*R*_, and Θ_*R*_ is the stimulus concentration that produces half-maximal response 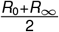. The dynamic range of the response is given by *R*_∞_ − *R*_0_, signifying the maximum the output can change in response to the input. (3) shows that shapes of dose response curves are same for the three signaling mechanisms. Hence comparison between them can be carried out in terms of *R*_0_, *R_∞_*, and Θ_*R*_.

At the receptor level, 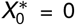 and 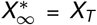, notwithstanding the rate parameters. The half-maximal stimulus Θ_*X*\*_ is equal to 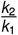, which is the binding affinity of the stimulus with the receptor. Furthermore, the fractional receptor occupancy 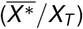 for a given stimulus (*S*) is determined by *k*_1_*S/k*_2_. As for the switch, the response 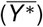 is specified by:

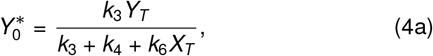

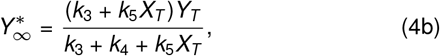

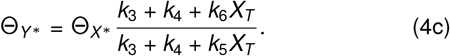

These expressions show that the dose-response of the switch depends upon the the basal rates as well as activation strength (*k*_5_*X_T_*) and repression strength (*k*_6_*X_T_*). A careful examination of (4) provides the following insights:

i. The activation strength (*k*_5_*X_T_*) does not affect the minimum response 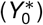, but affects the maximum response 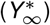. In particular, increasing *k*_5_*X_T_* increases 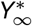. The repression strength (*k*_6_*X_T_*) decreases 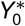 and does not affect 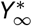.
ii. Relative values of the repression and activation strengths dictate the relationship between the half-maximal stimulus for the switch response (Θ_*Y*\*_) vis-á-vis the half-maximal stimulus for the receptor occupancy (Θ_*X*\*_). More specifically, Θ_*Y*\*_ < Θ_*X*\*_ when *k*_5_*X_T_* > *k*_6_*X_T_*, Θ_*Y*\*_ = Θ_*X*\*_ when *k*_5_*X_T_* = *k*_6_*X_T_*, and Θ_*Y*\*_ > Θ_*X*\*_ when *k*_5_*X_T_* < *k*_6_*X_T_*. Increasing *k*_6_*X_T_* increases Θ_*Y*\*_ while increasing *k*_5_*X_T_* does the opposite.

Fig. 2 illustrates the aforementioned effects on dose-response curves for the signaling mechanisms considered. Noting that signaling pathways typically show little activity in absence of the stimulus 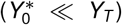 and show full activity 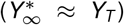 if the stimulus is large, it is reasonable to make the following assumptions: *k*_3_ ≪ *k*_4_ + *k*_6_*X_T_* and *k*_4_ ≪ *k*_3_ + *k*_5_*X_T_*. The limiting case of *k*_3_ = 0 leads to 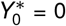; likewise, *k*_4_ = 0 results in 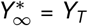. With these assumptions in mind, we use the following parameters for Fig. 2: *k*_3_ = 0 and *k*_6_ = 0 for activation; *k*_4_ = 0 and *k*_5_ = 0 for derepression; and *k*_3_ = 0 and *k*_4_ = 0 for concerted. As shown in Fig. 2(a) activation makes the switch response more sensitive to stimulus than the receptor occupancy (Θ_*Y*\*_ < Θ_*X*\*_). Increasing the activation strength (*k*_5_*X_T_*) increases 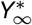 and decreases Θ_*Y*\*_, increasing the dynamic range (vertical expansion) and sensitivity (leftward shift) of the dose-response curve. The derepression mechanism exhibits an opposite behavior with Θ_*Y*\*_ > Θ_*X*\*_. In this scenario, increasing the repression strength increases the dynamic range by decreasing 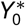 and decreases sensitivity by increasing Θ_*Y*\*_ (Fig. 2(b)).

**Figure 2:**
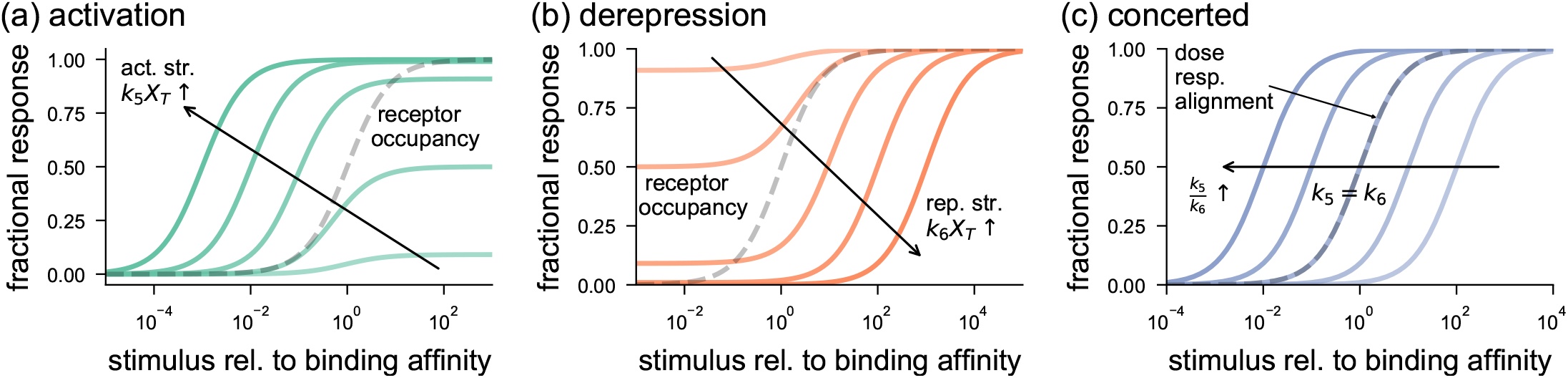
Dose-response curves for signaling mechanisms through molecular switches. The response is measured in terms of fraction of active switches *Y*\*/*Y_T_* as the stimulus level varies. The receptor occupancy curve denotes the fraction of active receptors *X*\*/*X_T_*. The stimulus is normalized by its binding affinity to the receptor (Θ_*X*\*_). **(a)** For the activation mechanism, the half-maximal stimulus (Θ_*Y*\*_) of a dose-response curve is less than Θ_*X*\*_. Each dose-response curve (solid line) is for a fixed activation strength *k*_5_ *X_T_*. Increasing *k*_5_ *X_T_*, depicted by the solid arrow, causes an upward expansion and leftward shift in dose-response. For these plots, the following values for parameters were used: *k*_3_ = 0, *k*_4_ = 1 and *k*_6_ = 0. The activation strength (*k*_5_ *X_T_*) was varied to take values from (0.1, 1, 10, 100, 1000). **(b)** For the derepression mechanism, the dose-response of the switch for a given repression strength (*k*_6_ *X_T_*) has half-maximal stimulus (Θ_*Y*_) greater than Θ_*X*\*_. Increasing *k*_6_ *X_T_*, shown by the solid arrow, leads to a downward expansion and rightward shift in the dose-response curve. The repression strength *k*_6_ *X_T_* takes values from (0.1, 1, 10, 100, 1000). The rest of the parameters were set as *k*_3_ = 1, *k*_4_ = 0, and *k*_5_ = 0. **(c)** In the case of concerted mechanism, Θ_*Y*\*_ may be greater than, equal to, or less than Θ_*X*\*_, depending upon the relative values of the activation strength and the derepression strength. Increasing the ratio *k*_5_ /*k*_6_, depicted by the solid arrow, shifts the dose response to left. Dose-response alignment (Θ_*Y*\*_ = Θ_*X*\*_) occurs when *k*_5_ = *k*_6_. The parameters used for the plots are *k*_3_ = 0 and *k*_4_ = 0. The ratio *k*_5_ /*k*_6_ was varied over (0.01, 0.1, 1, 10, 100).

Because we ignore the basal rates, changing activation and derepression strengths only influence Θ_*Y*\*_ in the case of a concerted mechanism. As expected, the switch response is more (less) sensitive than the receptor occupancy if activation (derepression) dominates derepression (activation). There is a perfect alignment of the fractional receptor occupancy curve with the dose response curve of the switch when *k*_5_ = *k*_6_ (Fig. 2(c)). Another important property of the concerted model is that it exhibits ratiometric signaling in which the response of the switch (*Y*\*) is determined by the ratio of active receptors to the total number of receptors (*X*\*/*X_T_*) [13, 14]. The absolute value of the total number of receptors (*X_T_*) has no bearing on 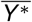. This may be seen by setting *k*_3_ = 0 and *k*_4_ = 0 in the expression of 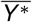 in (2):

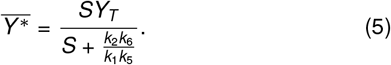

In reality, *k*_3_ and *k*_4_ are likely to be small, but non-zero. Therefore, ratiometric signaling does not hold in a strict sense.

Our theoretical results above show how the dose-response curves behave differently for activation, derepression, and concerted mechanisms. Are some of these behaviors observed in biological systems? One example where the signaling response becomes maximal when only a small fraction of receptors are bound (Θ_*Y*\*_ < Θ_*X*\*_) is the EGFR–MAPK pathway which elicits a full MAPK response at less than 5% receptor occupancy [57]. Our analysis explains this by an activation mechanism or a concerted mechanism in which the activation strength dominates the repression strength. A contrasting behavior is seen in the ethylene pathway of *Arabidopsis thaliana* in which a loss-of-function mutation of one of the ethylene receptors, *etr1*, shows increased sensitivity to etylene [58]. This points to a derepression mechanism in which the decreased amount of the receptor (*X_T_*) lowers the repression strength *k*_6_*X_T_* and shifts the dose response curve to the left in comparison to that of the wild-type system. A suggested example of concerted mechanism is the yeast G-signaling pathway, which exhibits both ratiometric signaling [13, 14] and dose-response alignment [43].

## Response times

Our analysis thus far focused on the steady-state properties of the activation, derepression, and concerted mechanisms. In this section, we study these mechanisms in terms of their response times; that is the time it takes for a signaling output to reach its steady-state. We use the following definition of response time:

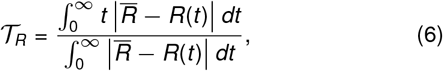

where *R*(*t*) is the time-dependent response of the pathway component under consideration and 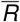 represents its value at steady-state [59]. For this definition, 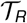 represents the “center of mass” of the response *R*(*t*), and is well-defined when 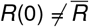. We may also think of 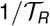 as the speed of the response in the sense that if the response is determined by a single kinetic step, 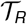 is reciprocal of the rate constant for that step. For example, the response time for the receptor is given by (section S1, SI):

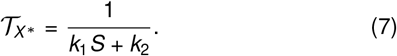

Thus, the response time decreases (i.e., response speeds up) if *k*_1_*S*+*k*_2_ increases. Because the response time depends upon the sum *k*_1_*S* + *k*_2_ and the steady-state receptor occupancy depends upon the ratio *k*_1_*S/k*_2_, these quantities can be tuned independently.

In the absence of stimulus, the response time of the switch follows the same form as (7):

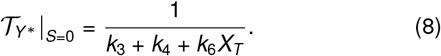

When the stimulus is present, analytic solutions to the integrals in (6) for the response time of *Y*\*(*t*) do not exist, except for a special case of the perfect concerted model *k*_5_ = *k*_6_. It is, however, possible to approximate 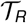 by linearizing the ODE system in (1) around its steady-state:

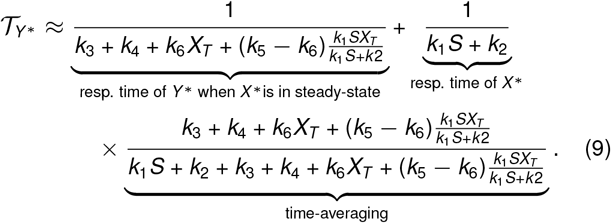

This equation is exact for the special case when *k*_5_ = *k*_6_ (section S2-B, SI). The first term in (9) can be interpreted as the response time of the switch when the receptors are at steady-state, because in that case the switch would be turned *on* at a rate 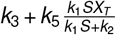 and turned *off* at a rate 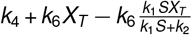; so, inverse of their sum would give the response time. The second term represents the response time of the receptor 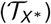 multiplied by a time-averaging factor which computes the ratio of 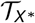 to the sum of 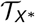 and the response time of the switch when 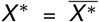. The time-averaging term lies between 0 and 1; its value approaches 0 if the receptor response is much faster than the switch response when 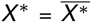 and approaches 1 if the receptor response is much slower than that of the switch.

If the receptor response is much faster than that of the switch, we expect that the latter does not depend upon the former (time-averaging term → 0). Indeed in this limit, (9) gives

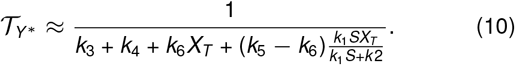

Comparing (10) with the basal response time in (8) shows that for a given stimulus level, activation speeds up the response in comparison with the basal response. In contrast, derepression slows down the response and a perfect concerted mechanism does not affect the response time (Fig. 3(a)). In the other limiting case when the receptor timescale is much slower than that of the switch, we expect the receptor dynamics to dictate the response time (time-averaging term → 1). Indeed in this case, (9) reduces to 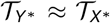 such that choice of the mechanism to control the switch has little effect on the response time. Our analytical as well as numerical calculations confirm this behavior (Fig. 3(a)).

**Figure 3:**
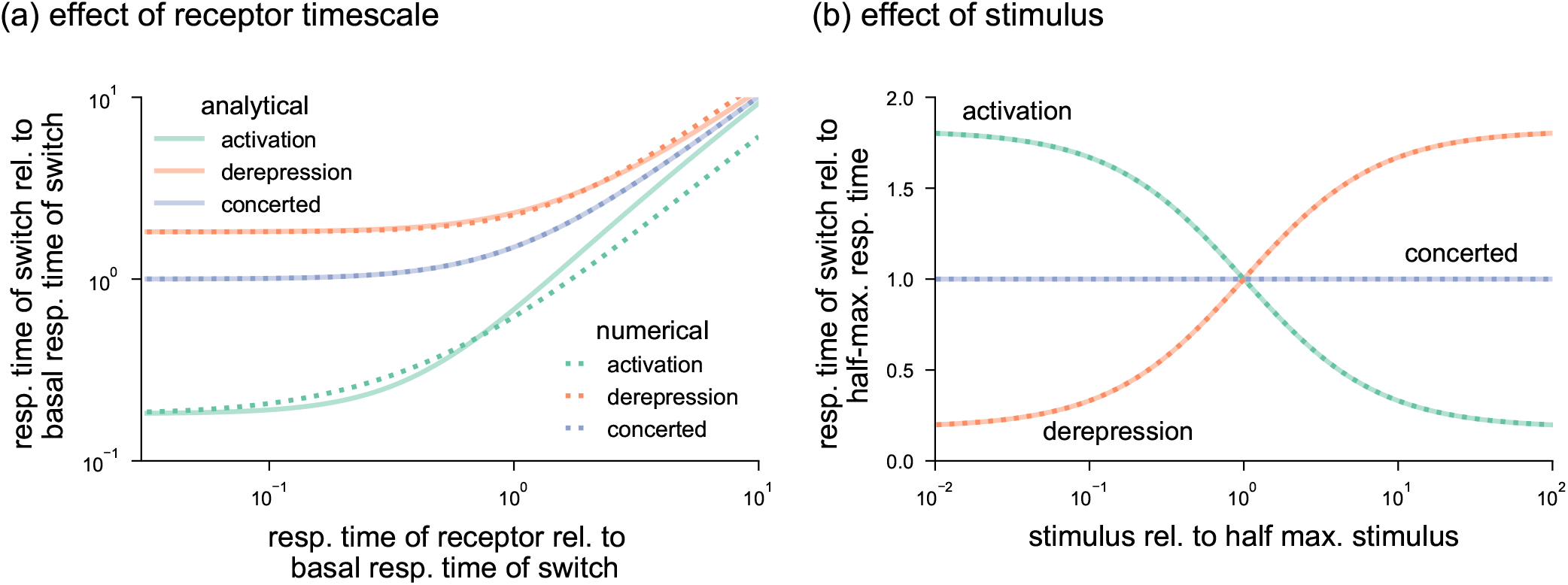
Response times of molecular switches governed by activation, derepression, and concerted mechanisms. (**a**) The response time of the switch increases as the response time of the receptor increases. Differences in the signaling mechanisms are more prominent when the receptor response is fast. Activation speeds up the response in comparison with the basal response time whereas dererepssion slows it down. A perfect concerted mechanism does not affect the response time. For each signaling mechanism, the response time is computed using the analytical result in (9) (solid lines) and numerically validated by using (6) (dashed lines). To ensure the same basal response and basal response time of the switches across signaling mechanisms, we chose the parameters as *k*_3_ = 1/9, *k*_4_ = 1, *k*_6_ = 0, and *k*_5_ *X_T_* = 10 for activation; *k*_3_ = 1/9, *k*_4_ = 0, *k*_5_ = 0, and *k*_6_ *X_T_* = 1 for derepression; and *k*_3_ = 1/9, *k*_4_ = 0 and *k*_5_ *X_T_* = *k*_6_ *X_T_* = 1 for concerted. The receptor response time was *k*_1_ *S* + *k*_2_ varied through *k*_2_ while maintaining *k*_1_ *S/k*_2_ = 1. (**b**) With increase in the stimulus level, response time decreases for activation, increases for derepression, and does not change for the concerted mechanism. The comparison is controlled by setting same response time at half-max. stimulus Θ_*Y*\*_. The following parameters were chosen to have same basal response but different basal response times: *k*_3_ = 1, *k*_4_ = 9, *k*_5_ *X_T_* = 90, and *k*_6_ = 0 for activation; *k*_3_ = 10,*k*_4_ = 0,*k*_5_ = 0, and *k*_6_ *X_T_* = 90 for derepression; and *k*_3_ = 10, *k*_4_ = 0, *k*_5_ *X_T_* = *k*_6_ *X_T_* = 90 for concerted. The receptor occupancy was varied by changing *k*_1_ *S/k*_2_ while maintaining the receptor response time 1/(*k*_1_ *S* + *k*_2_), which was chosen to be 100 times faster than the response time of the switches at their respective half maximal stimulus levels.

Next we examine the scenario where the switch is controlled by varying the stimulus level (*S*). Because changing the stimulus affects the response time of the receptor, which in turn affects the response time of the switch, we control for this effect by keeping *k*_1_*S* + *k*_2_ constant. We find that activation shortens the response time (speeds up the response) with increasing stimulus levels, whereas derepression increases the response time (slows down the response) (Fig. 3(b)). Importantly, the response time of the concerted mechanism is independent of the stimulus strength, and, therefore able to respond rapidly over the whole range of stimulus levels. To better understand this behavior, consider the response time for the limiting case of fast receptor dynamics. (10) can be rewritten as

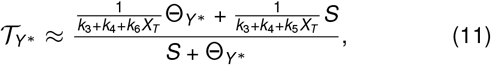

which changes from 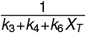 at *S* = 0 to 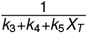 as *S* → ∞. The half-maximal stimulus Θ_*Y*\*_ is same as defined in (4). For the activation mechanism, 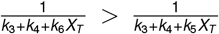; so the response time decreases with stimulus. Moreover, 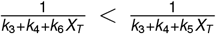 for the derepression mechanism and 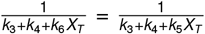 for the perfect concerted mechanism. Therefore, the response time increases with stimulus for the derepression mechanism and is independent of the stimulus for the concerted case.It is also worth pointing out that activation is faster than derepression only if the basal response times are equal. Therefore to construct a switch that responds rapidly using derepression, it is necessary for the switch to undergo fast basal cycling (Fig. 3(b)).

Our analysis of dose-response properties for ratiometric signaling given in (5) reveals that this mechanism is independent of the total number of receptors *X_T_* when the basal rates of the switch are zero (*k*_3_ = 0 and *k*_4_ = 0). Using these values in the expression for the response time in (9) demonstrates that this property does not hold for the response time. Specifically, the response time decreases with an increase in *X_T_* (section S2–B–c, SI).

Is there an intuitive explanation to why activation is faster than derepression? The activation model shortens the average life-time of the *off* state, without affecting the average lifetime of the *on* state. Derepression operates differently; it does not affect the average lifetime of the *off* state, but increases the lifetime of the *on* state. Thus, activation responds faster than derepression. The concerted mechanism simultaneously decreases the lifetime of the *off* state and increases the lifetime of the *on* state. Therefore, its response time lies between those of activation and derepression.

## Processing upstream fluctuations

The deterministic models used to compare the signaling mechanisms thus far ignore the stochastic nature of biochemical reactions, which becomes relevant when the abundance of receptor and switch proteins are small [48–53, 60, 61]. Therefore, we formulate a stochastic model of the concerted mechanism and analyze the other two mechanisms as its special cases. Our model consists of four reactions: activation of receptor upon recognizing the stimulus, deactivation of receptor, *on* to *off* transition of the molecular switch, and *off* to *on* transition of the molecular switch. The stochastic model is characterized by the probabilistic nature of each reaction and the discreteness of changes in population counts upon occurrence of a reaction as tabulated in Table 1.

**Table 1:**
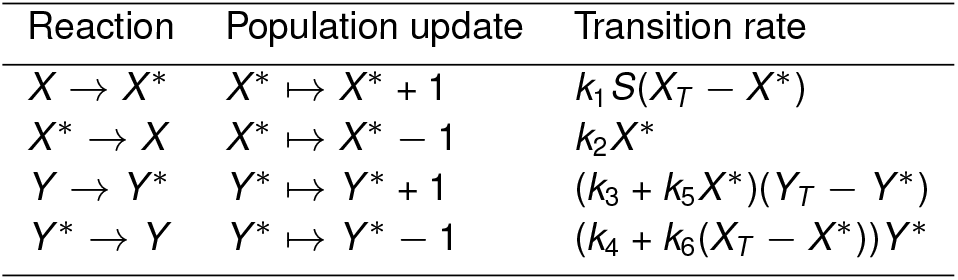
Transitions and associated rates for the stochastic model.

Our goal is to analyze the noise properties of activation, derepression, and concerted mechanisms. We quantify noise using coefficient of variation squared (*CV* ^2^), which is computed by normalizing the variance by mean^2^ and is a dimensionless quantity. To this end, we use the ODEs that describe the time evolution of the first and second-order moments, and solve them in steady-state to obtain the stationary moments [62–64] (section S3-B, SI). In particular, moments for the number of active receptors (*X*\*) are given by

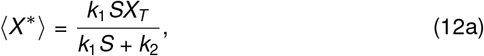

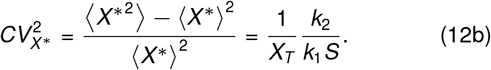

Here 〈.〉 denotes the expected value (average) of its argument. These moments correspond to a binomial distribution with parameters *X_T_* and 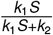 (section S3-A, SI). The stochastic mean 〈*X*\*〉 is same as the steady-state value for *X*\* in the deterministic model in (1). The coefficient of variation squared increases as the number of receptors (*X_T_*) decreases. Therefore the noise analysis is important when *X_T_* is small. In addition, the noise decreases with the ratio *k*_1_*S/k*_2_. Recall that *k*_1_*S/k*_2_ is the stimulus level relative to the binding affinity. Thus the noise diminishes when the stimulus level is much higher than the binding affinity.

Closed-form expressions for the moments are not available for *Y*\* owing to the nonlinear term *X*\**Y*\* in reaction rates, except for the special case of a perfect concerted model (*k*_5_ = *k*_6_). We approximate the mean response and the noise by considering a linearized system around the steady-state

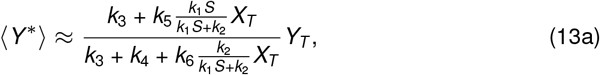

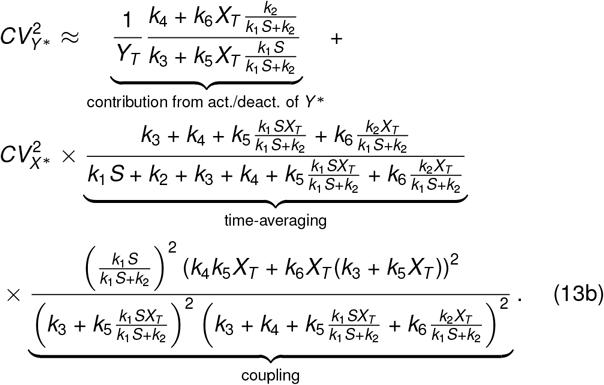

We validate these approximations using exact semi-analytical approach based on [65] (section S3-B, SI). The formula for 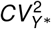 above is written in terms of various sources of noise, as previously done for gene regulation models [66–68]. Specifically, the noise in the signaling activity of the switch arises from two sources: activation/deactivation reactions of the switch, and noise in the number of active receptors (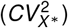. The contribution from activation/deactivation of the switch in (13) has a similar form as 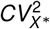 in (12). Accordingly, the contribution of this term decreases with increase in *Y_T_* or increase in the ratio of the total activation rate 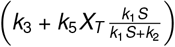 with total deactivation rate 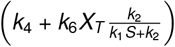. This ratio increases if the activation strength increases or the repression strength decreases. The contribution of 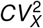 to 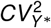 is scaled by time-averaging and coupling terms. The time-averaging term is the same as that in (9); it varies between 0 and 1, depending upon the relative timescales of the receptor and the switch. Thus, in the limiting case where receptor dynamics is very fast, the contribution from 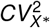 to 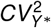 becomes negligible due to efficient time-averaging of fluctuations in *X*\*. The coupling term in (13) determines how strongly *X*\* affects *Y*\*. For example, this term is zero when the stimulus is absent (*S* = 0) or when both *k*_5_ and *k*_6_ are zero. In both these cases, the switch is decoupled from the receptor.

Next, we compare the noise properties of activation, derepression, and concerted mechanisms. To mathematically control the comparison, we assume that the receptor dynamics is same across the three strategies. In addition, we maintain the same average rate at which the switch turns *on* from the *off* state, i.e., 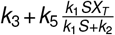, and the same average rate at which the switch turns *off* from the *on* state, i.e., 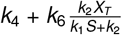. These assumptions ensure that differences in the noise properties, if any, are solely due to the architecture of the molecular switch and not dependent on the parameters. With this setup, we examine the effect of relative timescales (response times) of the receptor and the switch. We observe that in (13), varying *k*_1_*S* + *k*_2_ while maintaining *k*_1_*S/k*_2_ only affects the time-averaging term; all other terms are not affected. As shown in Fig. 4(a), the noise properties of these signaling mechanisms are similar when the receptor timescale is fast. This is expected because the dominant contribution in 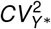 comes from its own activation and deactivation. However, when the receptor timescale is slower than that of the switch, the overall noise increases regardless of the signaling mechanism and the noise performance of the concerted mechanism becomes worse than the other two mechanisms.

**Figure 4:**
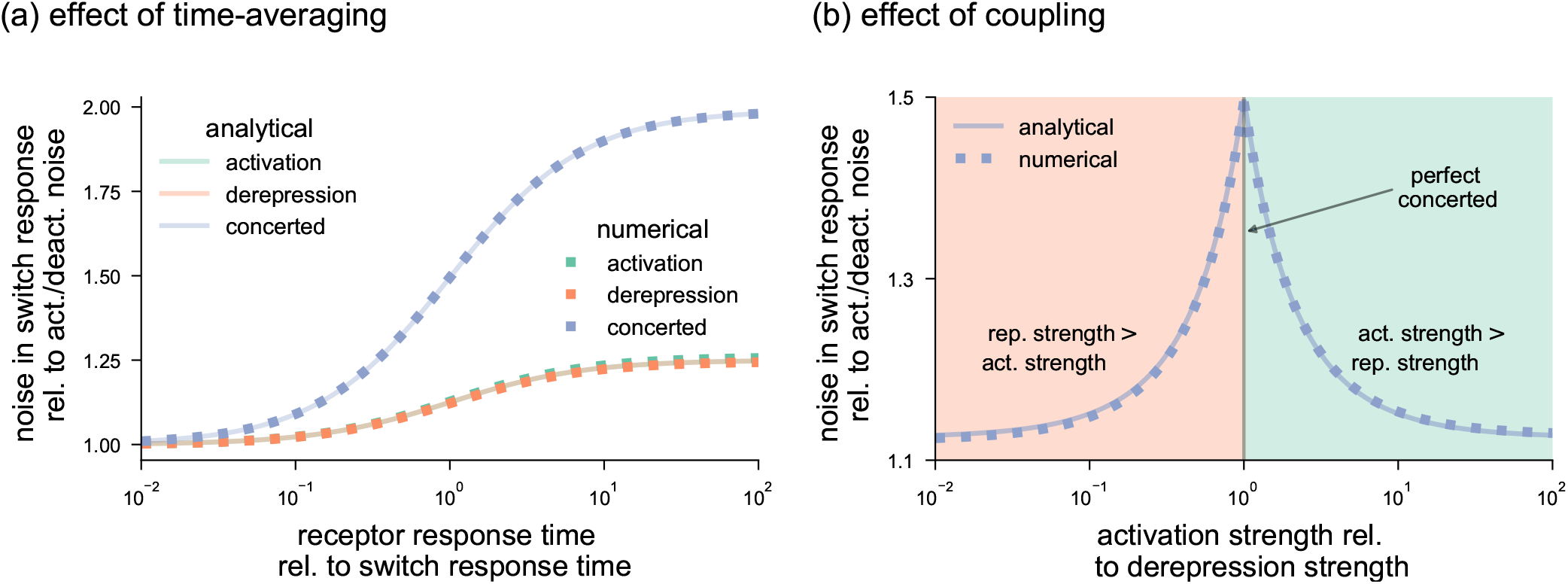
Noise in the number of active switch molecules. Noise is quantified using coefficient of variation squared 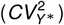 as in (13). The overall 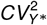 is shown relative to the contribution from activation/deactivation of *X*\*. The analytical result is computed using (13), which is validated numerically. (**a**) Noise with change in response time of the receptor. The noise increases as receptor response time increases, i.e., as the receptor slows down in comparison with the switch response time. The concerted model has a higher noise than activation and derepression, which perform similar. The difference is negligible when receptor dynamics is fast, and is more prominent when receptor is slow. The receptor response time (*k*_1_ *S* + *k*_2_) is varied by changing *k*_2_ while keeping the same receptor occupancy through the ratio *k*_1_ *S/k*_2_, so as to keep the same number of switches.The differences across signaling mechanisms are controlled by ensuring the same total activation rate of the switch 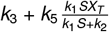 and same total deactivation rate 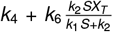. We used the following parameters: *k*_3_ = 0, *k*_4_ = 1, *k*_5_ = 0.02, and *k*_6_ = 0 for activation; *k*_3_ = 1, *k*_4_ = 0, *k*_5_ = 0, and *k*_6_ = 0.02 for derepression; and *k*_3_ = 0, *k*_4_ = 0, *k*_5_ = *k*_6_ = 0.02 for concerted. In addition, *X_T_* and *Y_T_* were taken to be 100 each. The receptor occupancy was maintained by *k*_1_ *S/k*_2_ = 1.(**b**) Noise with change in relative strengths of activation and derepression for a concerted mechanism. The noise is highest when the activation and derepression strengths match (perfect concerted mechanism). Deviating from the perfect concerted mechanism towards either stronger activation (shaded green region) or stronger derepression (shaded orange region) leads to smaller noise. Parameters were chosen such that total activation and the total deactivation rates were same across signaling mechanisms. For derepression, the activation strength was kept constant and the repression strength *k*_6_ *X_T_* was varied with a commensurate change in the basal deactivation rate *k*_4_. For activation, the repression strength was kept constant and the activation strength *k*_5_ *X_T_* was increased with appropriate change in the basal activation rate *k*_3_. We used the following parameters: *k*_1_ = 1, *S* = 1, *k*_2_ = 1, 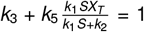, 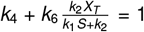, *X_T_* = 100, and *Y_T_* = 100.

The observation that activation and derepression both have similar noise and their concerted action has higher noise is surprising in light of our analyses of dose-response and response time. In terms of these properties, activation and derepression counteract to enable intermediate response for the concerted mechanism. Intuitively, the increase in fluctuations occurs because in the concerted mechanism, fluctuations in the upstream component affect both transitions *Y* → *Y*\* and *Y*\* → *Y*. In the case of activation and derepression, however, only one of these transitions is coupled with the upstream component. As a result, the concerted mechanism performs worse in terms of noise. We further highlight this observation by varying the relative strengths of activation (*k*_5_*X_T_*) and derepression (*k*_6_*X_T_*) in Fig. 4(b). The noise is greatest for the concerted mechanism when *k*_5_*X_T_* = *k*_6_*X_T_*.

We also analyze the special case of ratiometric signaling. Our deterministic analysis shows that for a concerted mechanism without basal rates (*k*_3_ = 0 and *k*_4_ = 0), the steady-state response 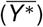 does not depend upon the total number of receptors (*X_T_*). However, similar to the response time, the 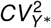 also depends upon *X_T_* through the time-averaging term and 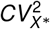, both of which decrease with increases in *X_T_* (section S3–B–c, SI). To summarize, ratiometric signaling only holds for the steady-state response. A cell that has higher *X_T_* would respond faster as well as with less noise than a cell with a smaller *X_T_*.

## Effect of receptor removal

Our models of signaling mechanisms in Fig. 1 assume conservation of number of receptor molecules (*X_T_*) and of switch molecules (*Y_T_*). These assumptions do not hold in case of some signaling pathways where stimulus-mediated removal of receptors occurs. Reported examples of such phenomena include GPCRs [54], EGFR [42], AMPA-type glutamate receptors [55], the receptor-like kinase FLS2 [56] and regulator of G-signaling (RGS) in *Arabidopsis thaliana* [11]. On the one hand, removal of active receptors is proposed to be a mechanism for desensitizing the response to a sustained stimulus [69, 70], and consequently enabling signaling over a broad range [41,42]. On the other hand, phosphorylation and subsequent removal of RGS, which is both a receptor candidate and a GAP, is proposed to result in sustained activation of signaling in *Arabidopsis thaliana* [9, 11, 45]. With an aim to explain these seemingly opposite behaviors of signaling pathways, we ask whether the signaling mechanisms, particularly activation and derepression, behave differently upon removal of receptors. To answer this, we reformulate the models in Fig. 1 by including production of inactive receptors (*X*) at a rate *k_p_*, removal of inactive receptors with rate *k_d_*, and removal of active receptors with rate 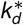. This model is simpler than those showing a broad range [41] or relative sensing [42], but is capable of adaptation [70] which is what we focus on.

Inclusion of receptor removal results in the following modification of the ODE system in (1)

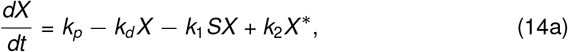

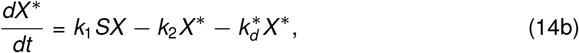

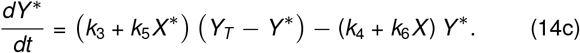

The initial conditions are: 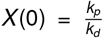, *X*\*(0) = 0, and 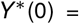 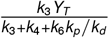. As before, setting *k*_6_ = 0 and *k*_5_ = 0, respectively, result in ODEs for the activation and derepression mechanisms.

An important distinction between the ODEs in (1) and the ODEs in (14) is that the receptor dynamics only has one timescale, 1/(*k*_1_*S* +*k*_2_), in the former but two timescales in the latter (section S4, SI). The interplay between these two timescales allows *X* (*t*) and *X*\*(*t*) to transiently respond to a stimulus at the fast timescale, followed by an eventual return towards their respective pre-stimulus levels at the slow timescale. Note that the switch response in the activation mechanism depends upon the active receptors *X*\*. Thus if *X*\* increases and returns towards its basal level, *Y*\* is also expected to follow the same dynamics. Likewise, if a derepression mechanism governs the switch then a decrease in *X* would lead to increase in *Y*\*. Further, if *X* returns towards its basal level, *Y*\* should also follow this trend. Such behavior is referred to as adaptation [70, 71].

What are appropriate parameter regimes where the switch response *Y*\*(*t*) in (14) adapts to a sustained stimulus? Our analysis shows that adaptation by *X*\* occurs when 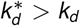, i.e., active receptors are removed at a faster rate than inactive receptors. In contrast, adaptation by *X* happens when 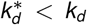 (section S4, SI). We note that all other parameters also affect the response properties, but the relative rates of receptor removal are the most important determinants of adaptive behavior. We illustrate these results in Fig. 5. To further bolster our observations, we examine the scenarios where the inactive receptors are preferentially removed for an activation mechanism and active receptors are preferentially removed for a derepression mechanism. Interestingly, in both these cases, the response sustains and does not adapt. These results thus provide another set of differences between activation and derepression mechanisms.

**Figure 5:**
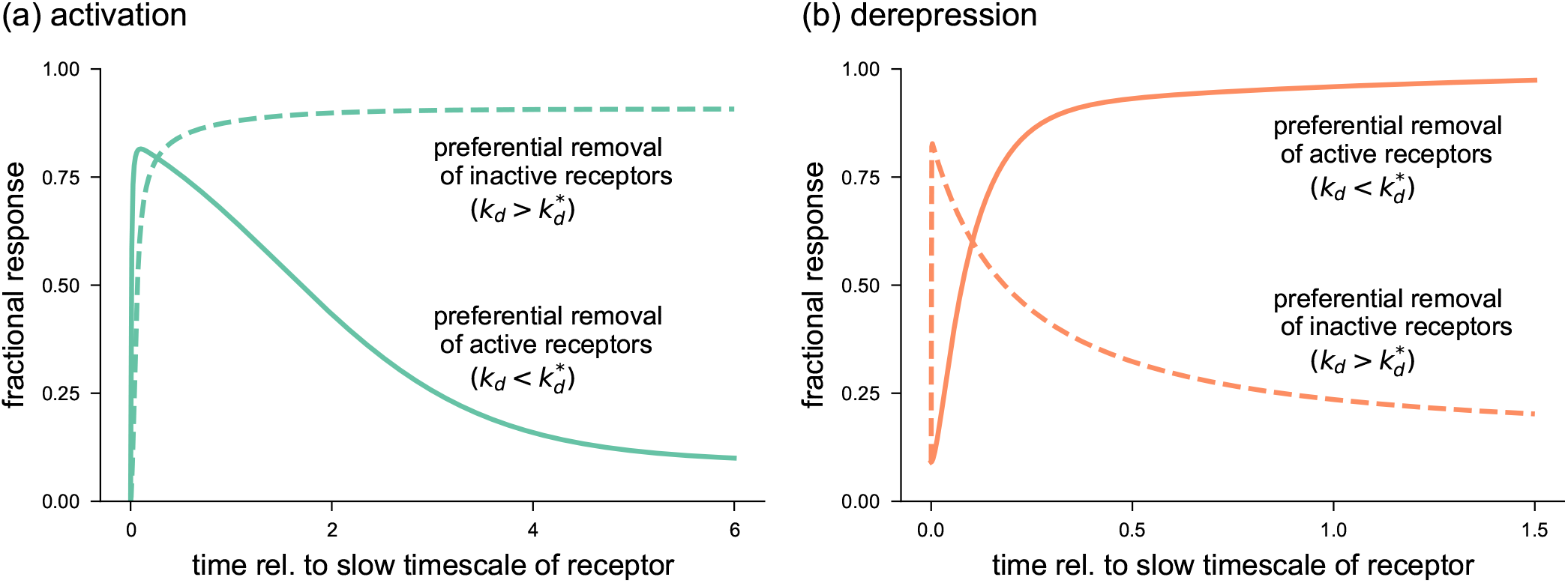
Effect of receptor removal on responses of activation and derepression mechanisms. The response is measured in terms of fraction of active receptors (*Y*\*/*Y_T_*) over time which is normalized to the slow timescale of the receptor (see section S4, SI). (**a**) For an activation mechanism, the switch response adapts, i.e., returns towards basal response after a transient, if the rate of removal of active receptors 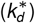 is higher than that of inactive receptors (*k_d_*). In contrast, if inactive receptors are removed at a faster rate, then the response sustains. For the adaptive response, we chose *k_p_* = 0.11, *k_d_* = 0.0011, 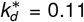. For the sustained response, we set *k_p_* = 101, *k_d_* = 1.01 and 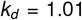. Rest of parameters were selected as *k*_1_ = 1, *k*_2_ = 1, *k*_3_ = 0, *k*_4_ = 10, *k*_5_ = 1, *k*_6_ = 0, *S* = 1, and *Y_T_* = 100 (**b**) For derepression mechanism, preferential removal of *X* results in adaptation whereas preferential removal of *X*\* causes sustained response.For the adaptive response, we used *k_p_* = 101, *k_d_* = 1.01, 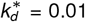. For the sustained response, we chose *k_p_* = 1, *k_d_* = 0.01 and 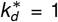. Rest of parameters were taken as *k*_1_ = 1, *k*_2_ = 1, *k*_3_ = 10, *k*_4_ = 0, *k*_5_ = 0, *k*_6_ = 1, *S* = 100, and *Y_T_* = 100

## Discussion

Molecular switches are important components of most signaling pathways. Typically, these switches can exist in two states, *on* and *off*, and the presence of an external stimulus biases the switch toward the *on* state. This transition can occur either by increasing the *off*-to-*on* rate (activation), decreasing the *on*-to-*off* rate (derepression), or both (concerted). We characterized these three mechanisms in terms of their dose-response curves, response times, and ability to process upstream fluctuations. We further examined how these three mechanisms were affected by receptor removal. The following list summarizes key differences in the performance of switches based on activation, derepression and concerted mechanisms:

- Both activation and derepression cannot align signaling activity with receptor occupancy. In particular, activation reduces the stimulus level required for half-maximal signaling as compared to 50% receptor occupancy (Θ_*Y*\*_ < Θ_*X*\*_), whereas derepression produces a rightward shift of the dose-response curve (Θ_*Y*\*_ > Θ_*X*\*_). The dose-response curve aligns with the receptor occupancy curve (Θ_*Y*\*_ = Θ_*X*\*_) for a perfect concerted mechanism (Fig. 2).
- A concerted mechanism is capable of ratiometric signaling, where the steady-state signaling output only depends upon fractional receptor occupancy and not on the total number of receptors.
- The response time for the activation mechanism decreases with signal strength, whereas it increases for the derepression mechanism. Importantly, the response time for a perfect concerted mechanism is independent of signal strength (Fig. 3).
- Activation and derepression mechanisms respond similarly to upstream fluctuations, whereas the concerted mechanism is more susceptible to fluctuations (Fig. 4). Unlike the mean steady state response, fluctuations in the output signal for the ratiometric signaling do depend on the total number of receptors.
- Preferential removal of active (inactive) receptors leads to an adaptive response for the activation (derepression) mechanism and a sustained response for the derepression (activation) mechanism (Fig. 5).

These results suggest performance trade-offs in the operating characteristics for each mechanism. The activation mechanism can increase the sensitivity of the pathway and generate response times that decrease with signal strength, but at the cost of dose-response curves that do not align with receptor occupancy, potentially limiting the pathways ability to transfer information [72]. In this sense, the activation mechanism operates as an ‘eager’ system that is sensitive to small receptor occupancies and accelerates the response for stronger signals. Therefore, activation seems appropriate for situations in which the cost of a false negative is greater than a false positive. For example, the adrenaline response to imminent danger should be sensitive and fast because cost of a false positive is small but a false negative can be deadly.

Similar to the activation mechanism, derepression leads to misalignment of the dose-response curve and receptor occupancy. However, for derepression the dose-response curve is shifted to the right. Another difference between these mechanisms is that for derepression, the response time increases with signal strength. Therefore, derepression acts as a ‘conservative’ system that does not respond to low receptor occupancy, waiting for a strong signal before committing to a response. Derepression seems appropriate for scenarios where the cost of a false positive is greater than a false negative. Interestingly, derepression-based signaling is found in many plants pathways. We speculate that it happens because plants have to continually allocate their limited resources between growth in competition with its neighbors and immunity to survive pathogen attack [73, 74]. For example, plants would perhaps ignore growth of a low level of pathogenic bacteria before allocating resources to fight them. Another possible scenarios where derepression may be used include irreversible cell-fate decisions such as the WNT pathway for embryo development [75], and fail-safe mechanisms such as the hypoxia-inducible factor in face of oxygen deprivation [76].

The concerted mechanism is better able to align with the receptor occupancy curve than either the activation or derepression mechanisms. Therefore, it has a better information fidelity [72]. The concerted mechanism also can generate response times that are independent of the strength of the input signal. However, these features come at the cost of higher susceptibility to upstream fluctuations. We note that in a recent study it was shown that ratiometric (concerted) signaling provided an advantage for gradient sensing, because it could compensate for spatial variations in the receptor concentration [14]. The system under consideration in that study was the mating response of yeast. For this case, the spatial fluctuations in the receptor concentration were larger than downstream fluctuations in signaling, allowing the concerted mechanism to outperform an activation-based mechanism.

While misalignment of the dose-response curve with receptor occupancy can cause loss of information, it may also offer some advantages. Consider a scenario where active receptors are preferentially removed, resulting in adaptation of the signaling response (Fig. 5). Recent work has shown that it is possible to exploit this feature to perform relative sensing (fold-change detection) if the receptor removal is a multi-step process [42]. Alternatively, a negative feedback may also result in an adaptive response and thereby a fold-change detection [59]. A key feature of fold-change detection is that the sensitivity of the system decreases each time the system adapts [59, 77]. Our results suggest that a relative sensing mechanism may be implemented with a derepression if the receptor removal operates on inactive receptors. We speculate that a negative feedback operating on inactive receptors would also yield the same effect.

Given that activation and derepression shift dose-response in opposite directions, a natural question to ask is whether dose-response alignment can occur in a signaling cascade where activation and derepression operate sequentially? To explore this possibility, we constructed a three-tier model where the response *Y*\* in Fig. 1(a) leads to derepression of a downstream component. Our analysis shows that indeed the response of the down-stream component is better aligned with the dose response than *Y*\*. We also analyze an alternate mechanism where derepression is followed by activation by modifying Fig. 1(b). As expected, the dose response of the downstream component aligns with the dose response better than that of *Y*\* (section S5, SI). It is worth noting that nonlinear regulation, such as feedback and feedforward loops, can also be used to compensate for undesirable characteristics of a given signaling mechanism. For example, negative feedback can align the dose-response curve with receptor occupancy for signaling pathways that operate through activation [43, 72, 78].

The models considered here are based on mass action kinetics and therefore cannot capture saturation effects. Traditionally, signaling pathways are modeled using Michelis-Menten kinetics. While we believe the qualitative features of our results will hold in this case, investigating how the behavior of the three mechanisms changes when the effects of enzyme saturation are included will be the subject of future work. Another future direction is to extend the analysis to include feedback and feedforward regulation. Finally, while we have focused our investigations on signaling pathways, our results are likely to be relevant in other intracellular systems, such as gene regulatory networks and metabolic pathways.

## Acknowledgments

The authors thank Daniel Lew (Duke University), Nicolas Buchler (North Carolina State University), Cesar A. Vargas-Garcia (Agrosavia), and members of the Jones and Elston labs for discussion and feedback. This work was supported by NSF grant MCB–1713880 to AMJ and TCE, NIH grant RO1 GM065989 to AMJ, and NIH grant R35 GM127145 to TCE.

## Supplementary Information

### S1 Comparison between different definitions of response time

Response time is a measure of the time it takes for a signaling output to reach its steady-state. In (6) of the main text, we defined the response time as the center of mass of the response curve. However, there are several interrelated definitions of response time. Here we provide a comparison between them. Towards that end, we use a model of a simple one-tier switch. Consider a protein that transitions between two states *A* and *A*\* as

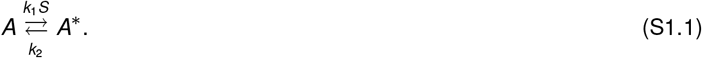

Let *A*(*t*) and *A*\*(*t*)denote the number of molecules that are in states *A* and *A*\*, respectively, at time *t*. We assume that the total number of molecules is conserved, i.e., *A_T_* = *A*(*t*) + *A*\*(*t*). We quantify the signaling through the switch by *A*\*(*t*), i.e., the number of molecules in the state *A*\*. The ordinary differential equation (ODE) governing the dynamics of *A*\* is:

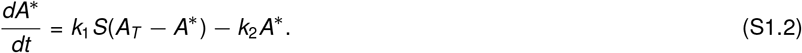

The solution to this ODE is given by

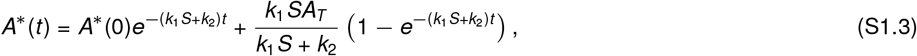

where *A*\*(0) < *A_T_* is the initial condition. As *t* → ∞, *A*\* approaches its steady-state value which is given by

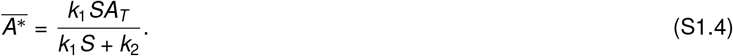

Recall the definition of response time from (6) in the main text

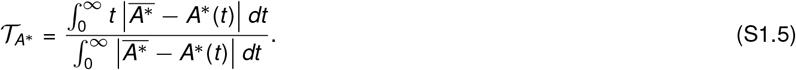

To compute the integrals in the numerator and the denominator, we first note that

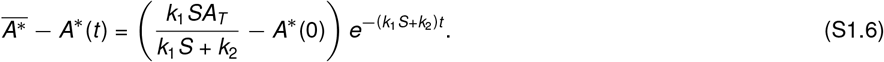

Because *k*_1_*S* > 0 and *k*_2_ > 0, 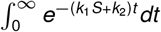 and 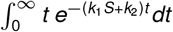 converge. These integrals are equal to 1/(*k*_1_*S* + *k*_2_) and 1/(*k*_1_*S* + *k*_2_)^2^, respectively. Using these integrals, (S1.5) gives

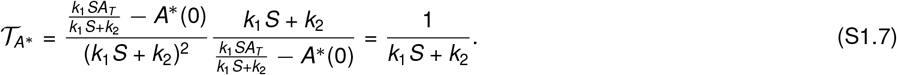

We can thus deduce that if the response is determined by a single kinetic step, the response time defined above is reciprocal of the rate constant for that step. It is also worth noting that the ratio is well-defined only when 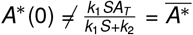.

Another class of definitions of response time are based on the time it takes for the response to start from *A*\*(0) and reduce its deviation from its steady-state by a factor 0 < *f* < 1. More specifically, we define 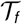 as the solution to the following equation

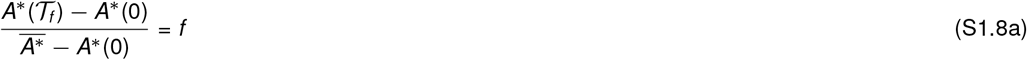

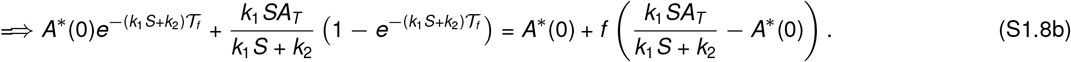

For 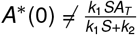, the above equation reduces to

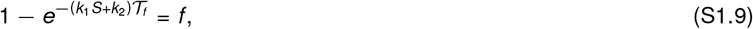

which has a straightforward solution

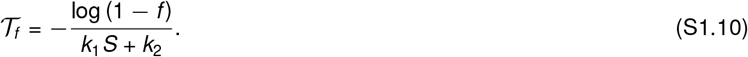

Notably, the response time is set by 1/(*k*_1_*S* + *k*_2_) up to a scale which depends on the specific value of *f*. We discuss three cases. First, setting *f* = 1/2 corresponds to the time at which half of the deviation from the steady-state has been reduced. The corresponding response time is given by

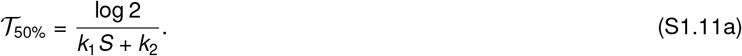

Second, *f* = (*e* − 1)/*e* ≈ 0.632 is also frequently used for which we obtain

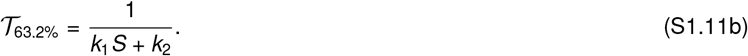

Lastly, a third definition concerns computing the time it takes for the response to travel from 10% to 90% of the difference between its initial value *A*\*(0) and steady-state 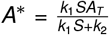. In this case, we get

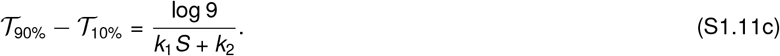

### S2 Transient solution and response time of two-tier cascades

In this section, we consider two-tier cascades of Fig. 1. Because activation and derepression are special cases of the concerted mechanism, we concern ourselves only with the ODEs of a concerted mechanism here.

The ordinary differential equations (ODEs) that govern the dynamics are

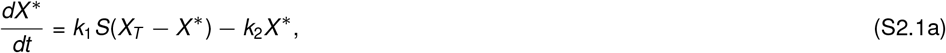

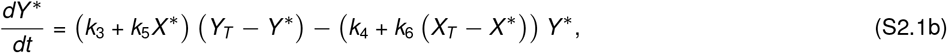

with initial conditions

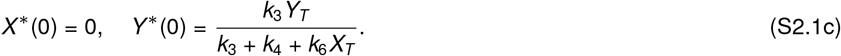

The steady-states of *X*\* and *Y*\* are computed by setting the derivatives to zero.

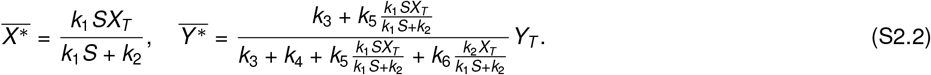

Recall that plugging *k*_6_ = 0 and *k*_5_ = 0, result in ODEs for the activation and derepression mechanisms, respectively. Furthermore, we term the special case *k*_5_ = *k*_6_ as perfect concerted mechanism, where the activation and repression strengths match.

#### S2-A Transient solution

Analytical solutions for nonlinear ODEs such as those in (S2.1) typically do not exist. However, a careful look at (S2.1) shows that the nonlinear term is (*k*_5_ *k*_6_)*X*\**Y*\*. Thus for a special case when *k*_5_ = *k*_6_ (perfect concerted mechanism), the system is linear, which exhibits analytical solution. The solutions for other cases can be computed numerically. We also provide an approximate solution using linearization around the steady-state solution 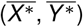.

It turns out that the forms of the ODEs for the perfect concerted mechanism and the linearized system are similar. Therefore, we consider the following generic system first and compute its transient solution.

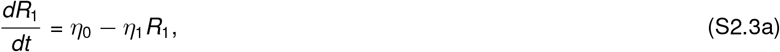

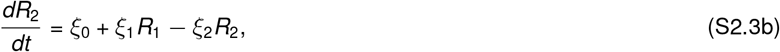

with initial conditions (*R*_1_(0), *R*_2_(0)). Let 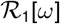 and 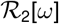 respectively denote the Laplace transforms of *R*_1_(*t*) and *R*_2_(*t*). Then we have that

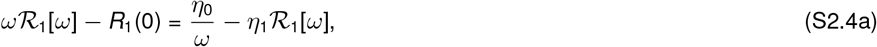

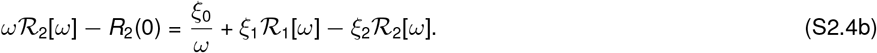

Solving for 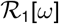 and 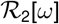

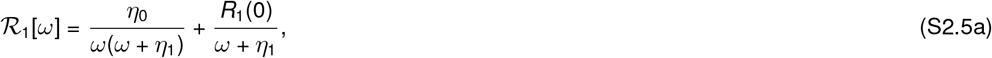

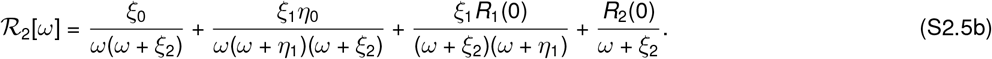

Taking inverse Laplace transform gives

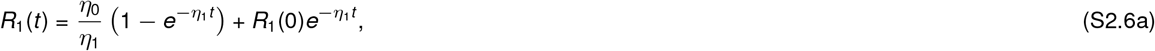

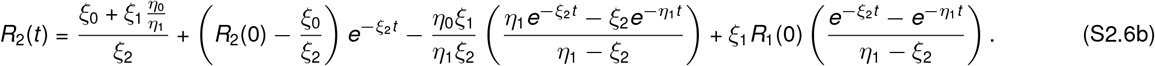

We can compute the steady-state solution by taking the limit *t* → ∞:

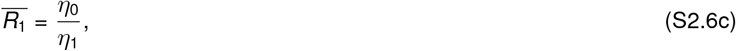

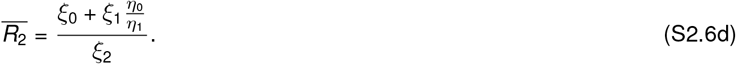

The solution for the limiting case when *η*_1_ = *ξ*_2_ may also be obtained by taking the limit *η*_1_ → *ξ*_2_. Another special case, which is more relevant for our discussion in this manuscript, is when the initial conditions are specified as *R*_1_(0) = 0 and 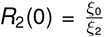. For this case, we have the following

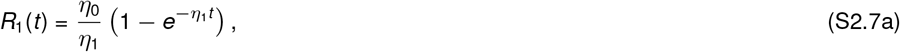

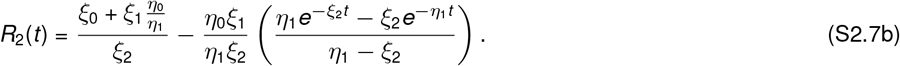

##### S2–A–a Transient solution for a perfect concerted model

A perfect concerted model is characterized by *k*_5_ = *k*_6_. Substituting *k*_5_ = *k*_6_ in (S2.1) results in

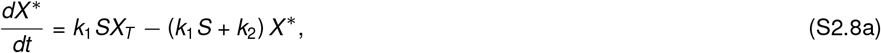

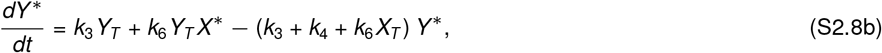

with initial condition 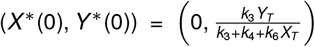. We note that the form of (S2.8) is same as that of (S2.3), with parameters *η*_0_ = *k*_1_*SX_T_*, *η*_1_ = *k*_1_*S* + *k*_2_, *ξ*_0_ = *k*_3_*Y_T_*, *ξ*_1_ = *k*_6_*Y_T_*, and *ξ*_2_ = *k*_3_ + *k*_4_ + *k*_6_*X_T_*. Thus, we can use (S2.7) to get the transient solution

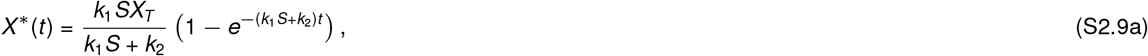

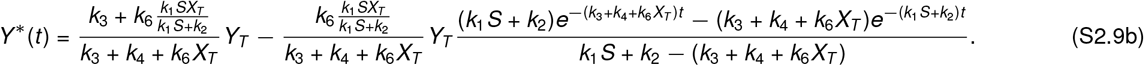

For the special case when *k*_1_*S* + *k*_2_ = *k*_3_ + *k*_4_ + *k*_6_*X_T_*, we have

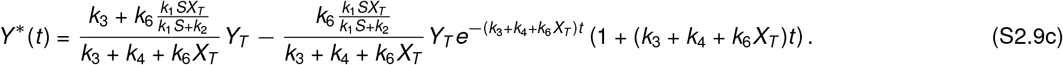

##### S2–A–b Approximate transient solution using linearization

The ODE system in (S2.1) contains the nonlinear term *X*\**Y*\*, which can be linearized around the steady-state solution 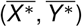 as

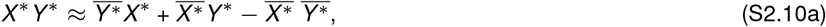

where

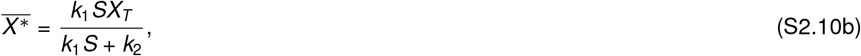

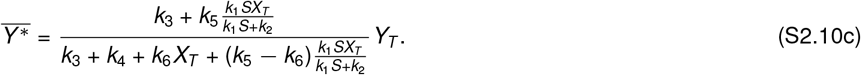

Substituting this for the nonlinear term in (S2.1), we get the following

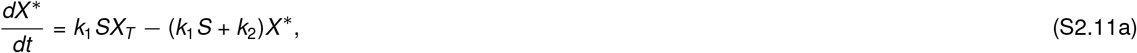

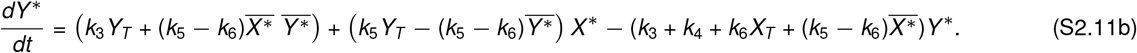

These ODEs are similar to those in (S2.3). The parameters are: *η*_0_ = *k*_1_*SX_T_*, *η*_1_ = *k*_1_*S* + *k*_2_, 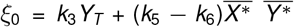, 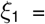 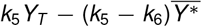, and 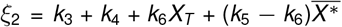. With the initial conditions 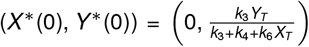, the solution same as that in (S2.7) and is given by.

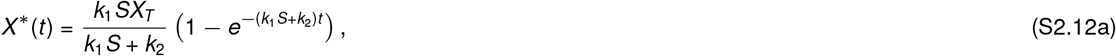

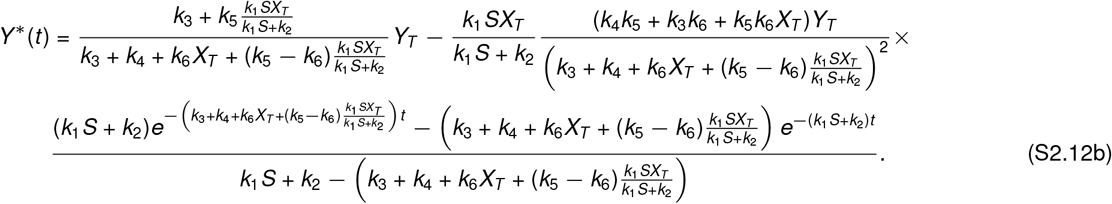

The special case when the timescales match may be computed by taking the limit of the above solution.

#### S2-B Response time

In this section, we compute the response times for the perfect concerted model and the linearized model. To this end, we recall that the response time for a response *R*(*t*) is

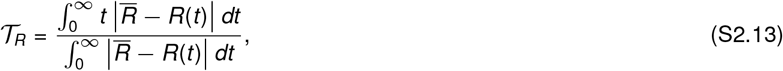

where 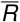 is the steady-state response. We use this definition to compute the response times for the generic ODE system considered in (S2.3), whose solution is given by (S2.6). We then adapt the solution for our systems of interest, namely, the perfect concerted model and the linearized model.

We begin by computing the response time for *R*_1_(*t*). The term 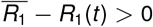 is

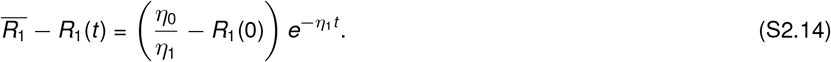

Note that

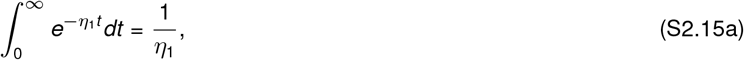

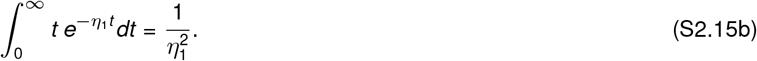

Using these in (S2.13), we get

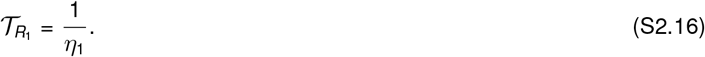

It is worth noting that 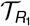 does not depend upon the initial condition *R*_1_(0) and is only defined if 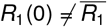.

Next we compute the response time for *R*_2_(*t*). The term 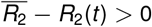 is given by

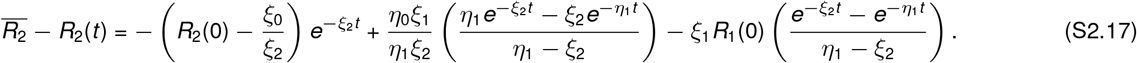

The integrals of exponential terms in (S2.15) may be used to compute the integrals for the numerator and the denominator of the response time. In particular, we have that

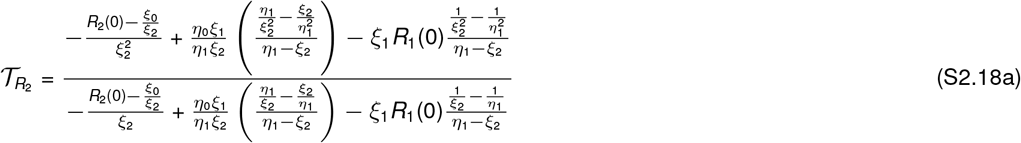

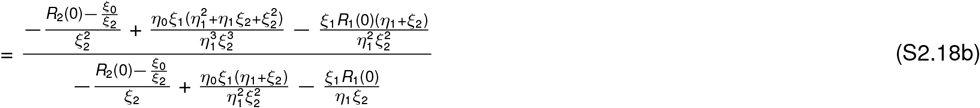

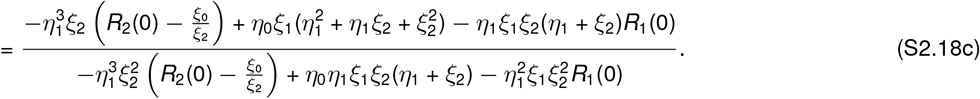

We deduce several important insights from the above expression. First, we note that the response time 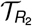 depends upon the initial conditions *R*_1_(0) and *R*_2_(0). Second, the dependence on *R*_1_(0) and *R*_2_(0) drops for the special case when *R*_1_(0) = 0 and 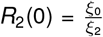. In this case, 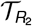 simplifies to

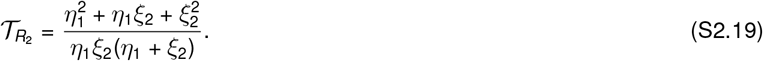

Finally, if *R*_1_(0) is taken to be at the steady-state 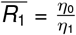 and *R*_2_(0) is set as 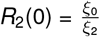, then we get

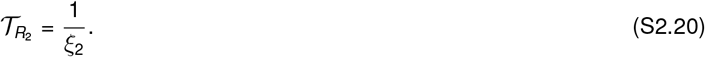

With this in mind, it is convenient to express (S2.19) as

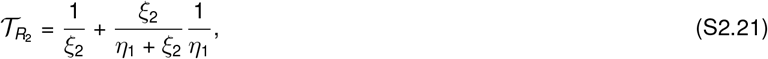

where the first-term is the response time if *R*_1_ were at steady-state, and the second term is the time-averaged 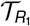.

##### S2–B–a Response time for a perfect concerted mechanism

For this case, we can simply adapt the results of (S2.16) and (S2.21).

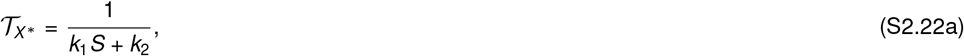

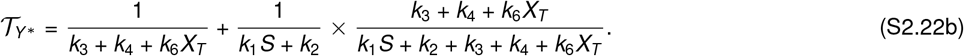

##### S2–B–b Response time for the linear approximation

As with the response time for the perfect concerted mechanism, here too we adapt the results of (S2.16) and (S2.21).

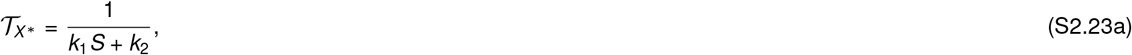

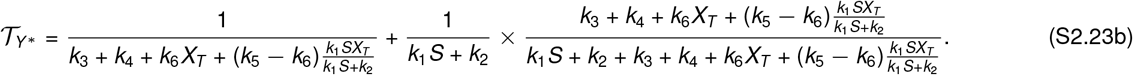

How good is the above approximation of response time? One check is to plug in *k*_5_ = *k*_6_ to obtain the approximation for the perfect concerted model for which we have the exact expression of the response time in (S2.22). Indeed, substituting *k*_5_ = *k*_6_ in (S2.23) yields

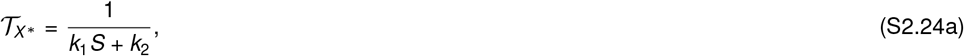

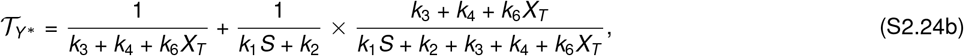

which is exactly same as (S2.22). Thus the linear approximation is exact for the perfect concerted model. This is not surprising because the perfect concerted model is linear by construction.A second check of how good the approximation in (S2.23) is through numerical computation, which is discussed in a later section.

##### S2–B–c Response time for ratiometric signaling

Ratiometric signaling is the special case where the signaling output does not depend upon the total number of receptors *X_T_*. In the main text, we show that when *k*_3_ = 0 and *k*_4_ = 0, then the response is independent of *X_T_* ((5)). Here we ask whether setting *k*_3_ = 0 and *k*_4_ = 0 also result in the response time indepedent from *X_T_*. To this end, we plug these values in the expression of 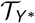 in (S2.23):

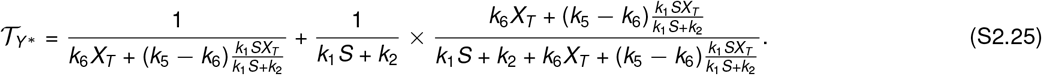

Clearly, the response time depends upon *X_T_*, thereby establishing that the ratiometric signaling is only applicable for the dose-response. We further ask how *X_T_* affects the response time. To this end, the most convenient limit to check is when the receptor dynamics is fast, i.e., 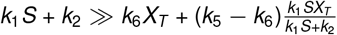, which gives us

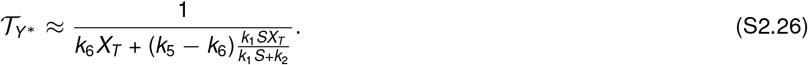

Thus, if everything else is constant then increasing *X_T_* decreases the response time. Even when the receptor dynamics is not fast, we can verify this effect by looking at the sign of the derivative of 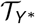 with respect to *X*\*

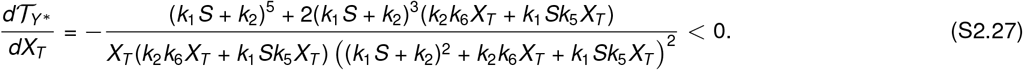

Thus increasing *X_T_* speeds up the response. Next, we discuss the numerical method to compute response time which we use to validate our approximations.

##### S2–B–d Numerical computation of the response time

One convenience in using the center of mass definition of the response time

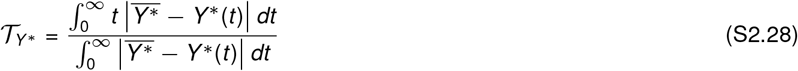

is that it can be computer numerically via solution of an augmented ODE system

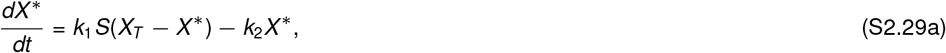

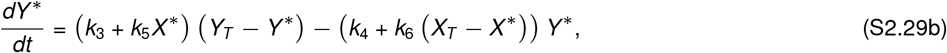

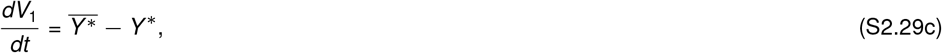

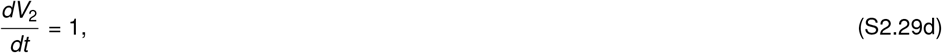

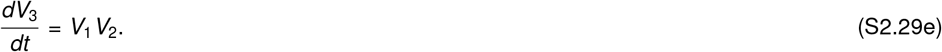

Here *V*_1_(*t*), *V*_2_(*t*) and *V*_3_(*t*) are the augmented states to the original ODE system. The initial conditions are given by

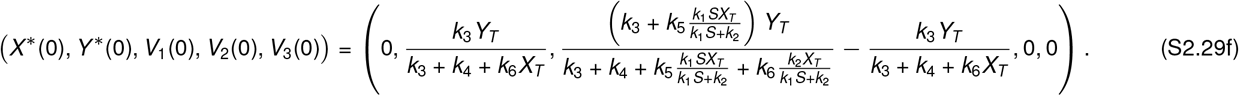

Note that the state *V*_1_(*t*) computes the integral in the denominator upto a time horizon *t*, *V*_2_(*t*) tracks the time, and *V*_3_(*t*) computes the numerator up to time horizon *t*. If we choose *t* to be large enough such that the system has reached saturation, then 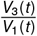 computes the response time. It is easy to see that the approximation gets better with a larger *t*. We can use the approximation of response time in (S2.23) to set a time for the integration.

### S3 Stochastic analysis of two-tier cascades

Here we consider a two-tier model for signal transduction as described in Table 1 in the main text. Let *P_m_*_,*n*_(*t*) denote the probability of finding *m* molecules of *X*\* and *n* molecules of *Y*\* at time *t*. Then, we can write the chemical master equation (CME) that describes the time evolution of *P_m,n_*

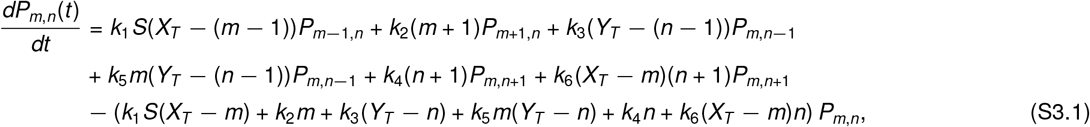

where *m* = 0, 1,…, *X_T_* and *n* = 0,…, *Y_T_* [60, 79]. It is often difficult to analytically solve the CME. Because the dynamics of *X*\* is linear and it does not depend upon *Y*\*, it is possible to provide an analytical solution *P_m_*. As for *P_m,n_*, we only provide approximate and exact computations of its first two moments.

#### S3-A Stochastic solution to receptor dynamics

The CME that governs the time evolution of *P_m_*(*t*) is:

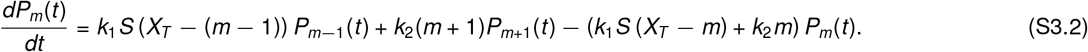

We define a generating function

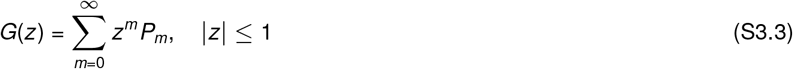

to solve (S3.2). Multiplying both sides by *z^m^* and summing over *m* yields

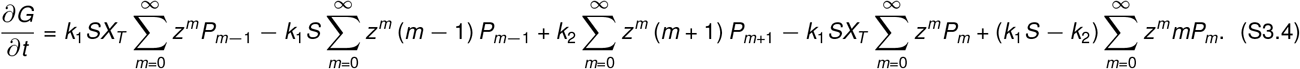

The above equation becomes the following partial differential equation (PDE)

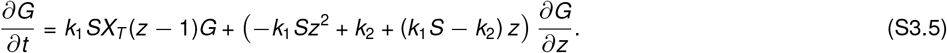

We solve this PDE using method of characteristics, assuming the initial condition *G*(*z*, 0) = 1 which corresponds to 0 molecules of *X*\*. The solution is given by

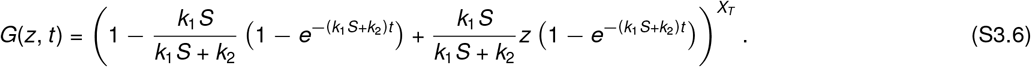

Using Binomial theorem, the above expression can be written as

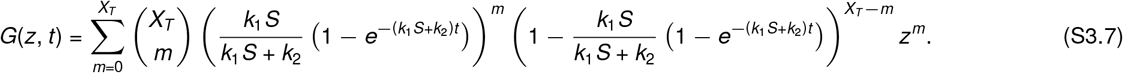

The probability *P_m_*(*t*) is given by the coefficient of *z^m^*

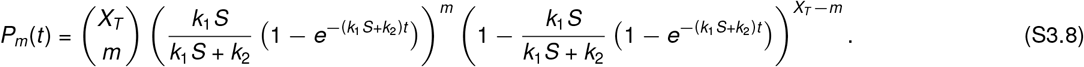

The stationary distribution 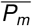 is computed by taking limit *t* → ∞

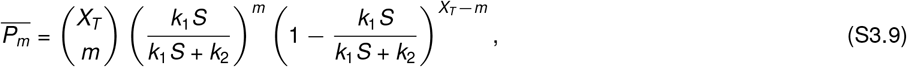

which is a Binomial distribution with parameters *X_T_* and 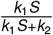 [80]. The stationary moments of this distribution are given by

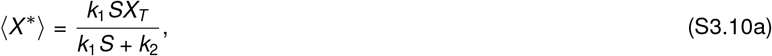

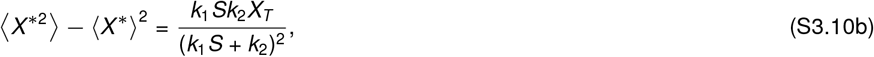

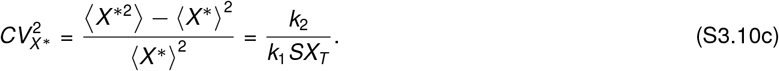

#### S3-B Moment dynamics

We are specifically concerned with moments of the two-tier model. To this end, we take the well-established approach of using the ODEs that govern the moment dynamics (e.g., see [62, 64]). A generic moment may be written as

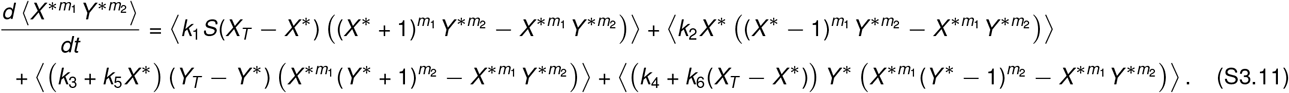

Here we have used 〈.〉 to denote the expected value of a random variable. Our focus in this work is to compute the first two moments in steady-state. However, due to the nonlinearity *X*\**Y*\* in these equations, the moment dynamics is not closed in that a lower-order moment depends upon a higher-order moment [62–64]. It turns out that for the special case *k*_5_ = *k*_6_ (perfect concerted model), the moments may be computed exactly. We provide approximate formulas for moments using a linear approximation when *k*_5_ ≠ *k*_6_.

##### S3–B–a Moment computation for a perfect concerted model

For the concerted model, *k*_5_ = *k*_6_. Let us write moment dynamics for first two moments.

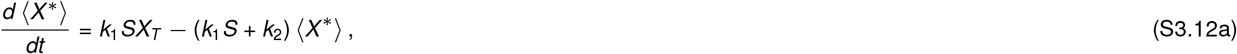

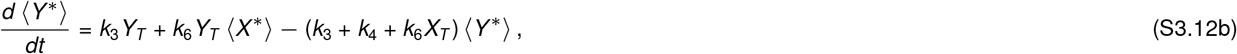

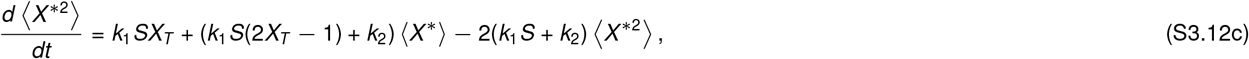

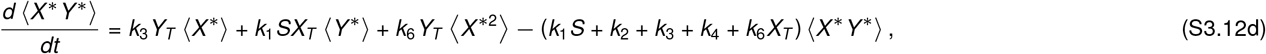

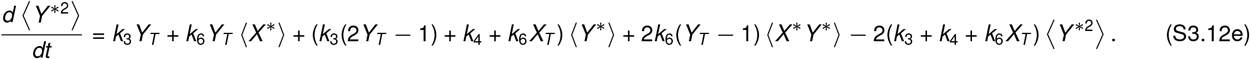

We can solve for steady-state moments by setting each of the derivatives equal to zero. For example, the means are given by

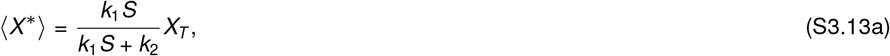

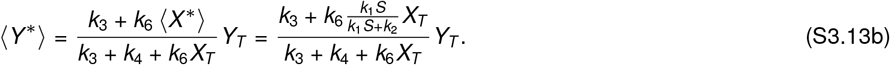

Next, we compute second order moments. 〈*X*\*^2^〉 is given by

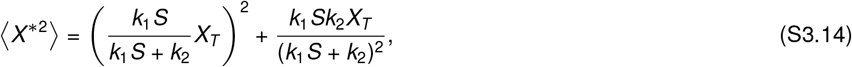

where the first term is 〈*X*\*〉^2^. The cross moment 〈*X*\**Y*\*〉 is

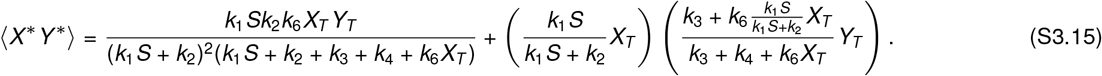

Here the second term is 〈*X*\*〉 〈*Y*\*〉. Finally, the second order moment 〈*Y*\*^2^〉 in terms of the other moments is

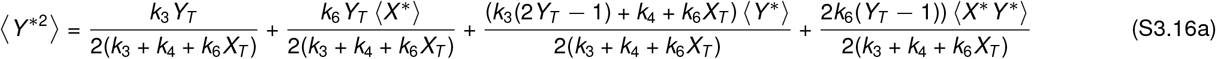

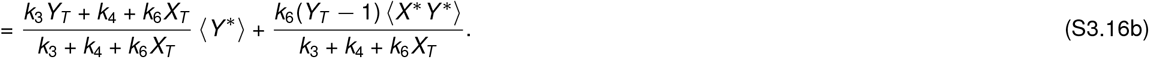

Using the moments computed above, we can compute the centered moments. For example, the variance of *X*\* is

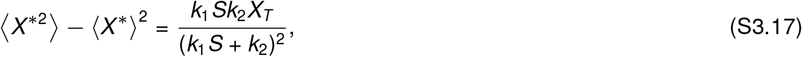

the centered cross moment is

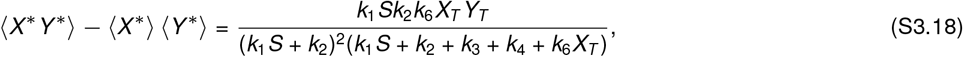

and the variance of *Y*\* is

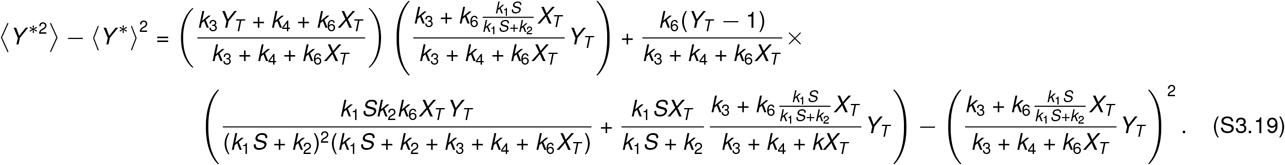

We use the centered moments computed above to quantify noise in *X*\* and *Y*\* using coefficient of variation squared.

###### Coefficient of variation squared

Let 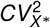 and 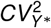 respectively are the coefficient of variation squared for *X*\* and *Y*\*. Then

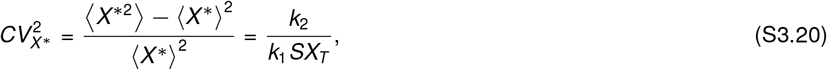

and

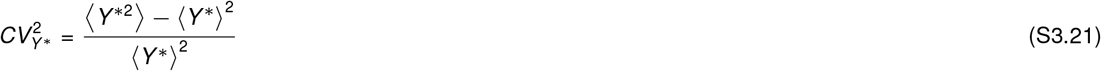

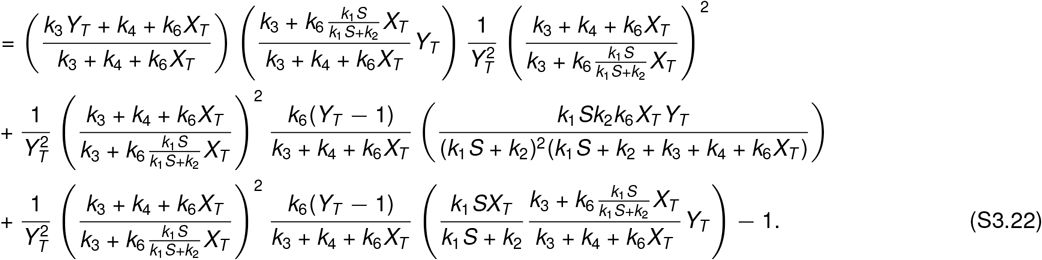

On simplifying, we get

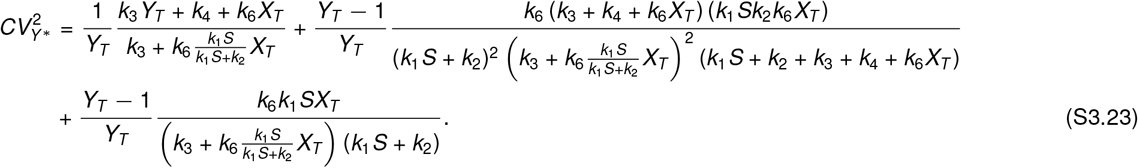

###### Decomposing the coefficient of variation squared into different sources

We expect that 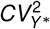 has two sources of noise: activation/deactivation events for *X*\* and activation/deactivation events for *Y*\*. To tease out the contribution from activation/deactivation events for *Y*\*, we consider a scenario the dynamics of *X*\* is deterministic. In this case, the moment dynamics is given by

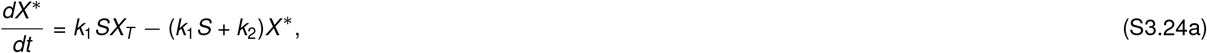

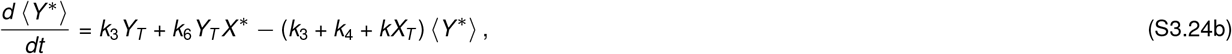

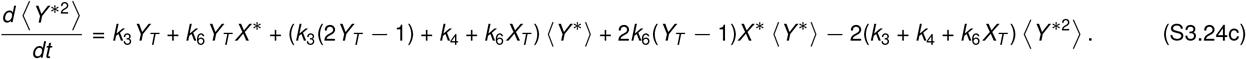

The steady-state solution for the coefficient of variation squared computed from these equations is given by

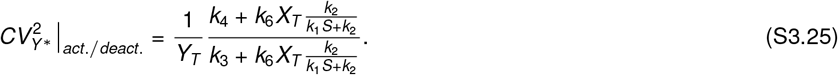

We do not provide detailed calculations here. One sanity check is that this expression is consistent with coefficient of variation squared for a binomial distribution, which is expected if *X*\* were constant.

Subtracting (S3.25) from (S3.23), we obtain the contribution of noise in *X*\* to noise in *Y*\*:

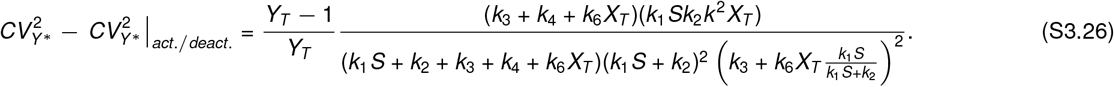

We expect that the term on the right hand side should have contribution from 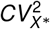, which is time-averaged. Recall (S2.22) that *k*_1_*S* + *k*_2_ is response time of the receptor and that *k*_3_ + *k*_4_ + *k*_6_*X_T_* is response time of the switch if the receptor dynamics is fast. Thus, 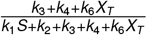 can be interpreted as the timescale averaging. Therefore, we write

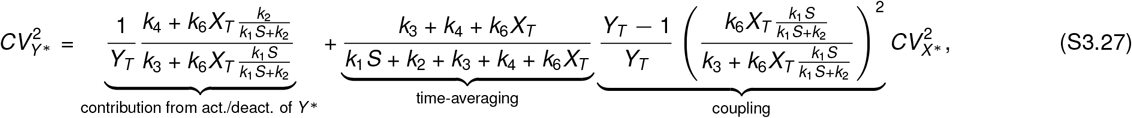

where 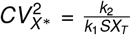.

##### S3–B–b Approximate moment dynamics using linear approximation

As discussed earlier, the moment dynamics is not closed when *k*_5_ − *k*_6_ is non-zero. To estimate moments, we first linearize the nonlinear term around the solution of the deterministic model [81]. Let 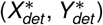 be solution to the ODE model

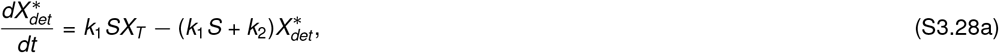

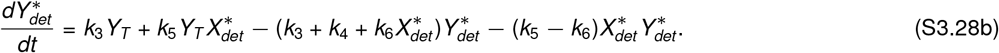

The stochastic model with linearized propensity is shown in Table 2.

**Table 2:**
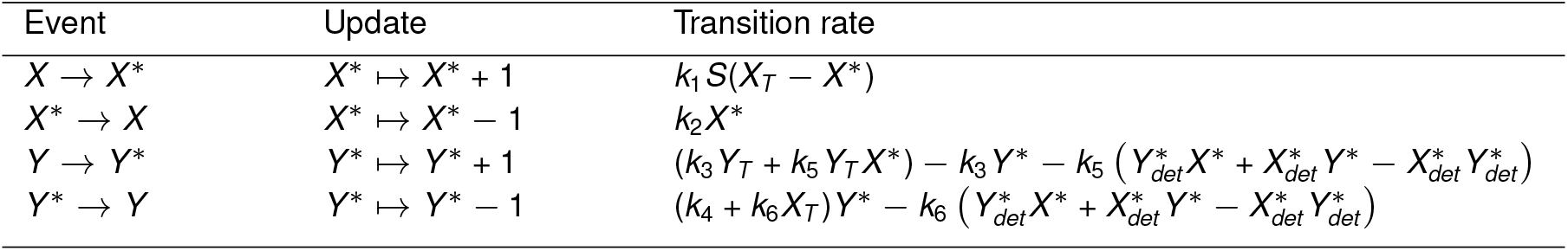
Transitions and associated rates for the stochastic model.

| Event | Update | Transition rate |
| --- | --- | --- |
| $X \rightarrow X^*$ | $X^* \mapsto X^* + 1$ | $k_1 S(X_T - X^*)$ |
| $X^* \rightarrow X$ | $X^* \mapsto X^* - 1$ | $k_2 X^*$ |
| $Y \rightarrow Y^*$ | $Y^* \mapsto Y^* + 1$ | $(k_3 Y_T + k_5 Y_T X^*) - k_3 Y^* - k_5 (Y_{det}^* X^* + X_{det}^* Y^* - X_{det}^* Y_{det}^*)$ |
| $Y^* \rightarrow Y$ | $Y^* \mapsto Y^* - 1$ | $(k_4 + k_6 X_T) Y^* - k_6 (Y_{det}^* X^* + X_{det}^* Y^* - X_{det}^* Y_{det}^*)$ |

The second order moments with the above linearized propensity model satisfy the following differential equations

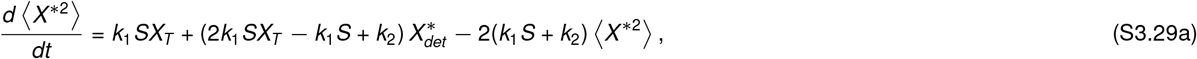

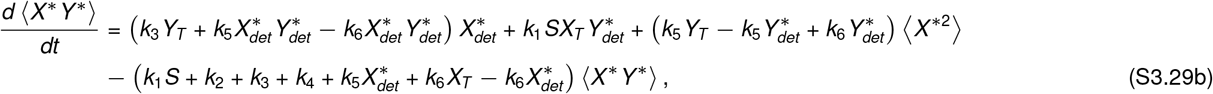

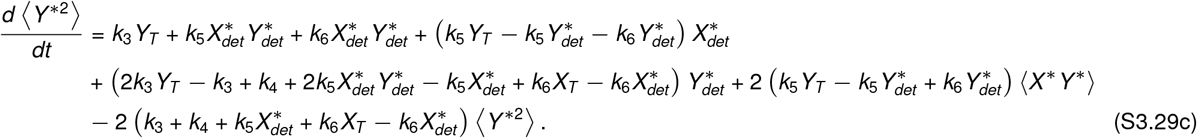

Computing these moment equations, along with the solutions to the deterministic dynamics, approximates the moments. Using a symbolic solver to solve for moments in steady-state, we get the following for the coefficient of variation of *X*\*.

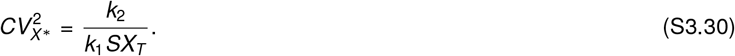

The formula for 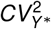 can be obtained in the same manner as done for the perfect concerted model and is given by

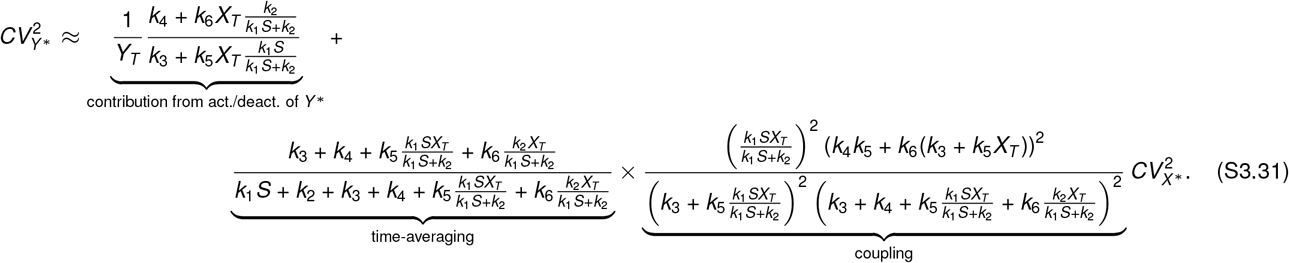

Because we already have exact moment formulas when *k*_5_ = *k*_6_, we can immediately check the validity of linear approximation for that case. Plugging *k*_5_ = *k*_6_ shows that the noise approximation above differs from (S3.27) by a factor (*Y_T_* − 1)/*Y_T_* that multiplies 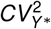. Typically (*Y_T_* − 1)/*Y_T_* ≈ 1 for large *Y_T_*, indicating that our linear approximation is reasonably good for a concerted model.

##### S3–B–c Coefficient variation squared for ratiometric signaling

For ratiometric signaling, in which the steady-state response does not depend upon the total number of receptors *X_T_*, we need *k*_3_ = 0 and *k*_4_ = 0. Substituting these in the expression of 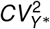 in (S3.31), we get

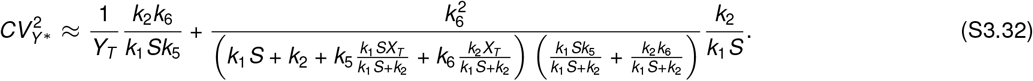

Thus, increasing *X_T_* decreases overall noise because *X_T_* increases the denominator terms in the above above formula. Next, we provide exact computation of moments using a semi-analytical approach.

##### S3–B–d Exact moment computation

Our goal here is to compute the first two moments of *Y*\*. As discussed earlier, a moment of lower order depends upon moments of higher order, resulting in the problem of moment closure. Here, we exploit the fact that *X_T_* is finite to come up with an alternate state space where moment dynamics is closed. The computations follow the formalism proposed in [65]. Another closely related method is the method of conditional moments described in [82].

Let us define indicator variables *b_i_*, *i* = 0, 1,…, *X_T_* as

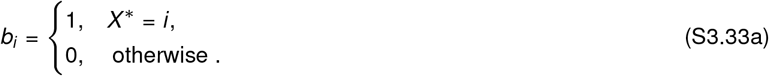

It then follows that

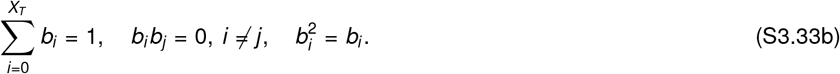

We now recast our original model in the new state-space 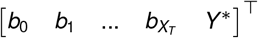. The transitions (i.e., reactions) and the corresponding transition intensities are as follows.

1. Receptor activation: the transition intensity of a receptor activation event is given by 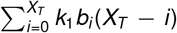. Whenever this event occurs, the states reset as

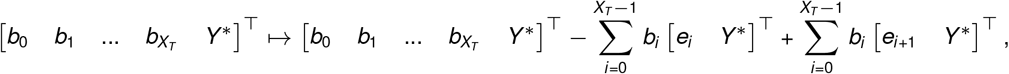

where *e_i_* is a column vector of dimension *X_T_* + 1, with all zeros except at the *i^th^* position. This reset map simplifies to

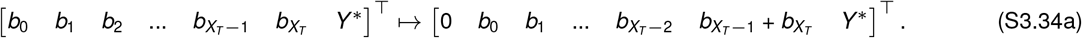
2. Receptor deactivation: the transition intensity is given by 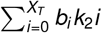, with the map

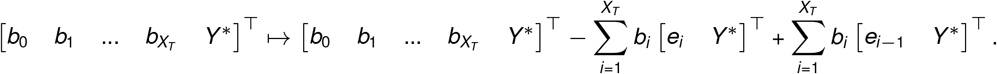

The reset map further simplifies to

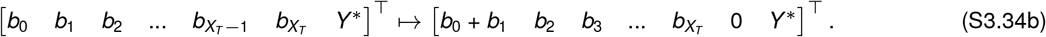
3. State *Y*\* to *Y*\* + 1 occurs with transition intensity 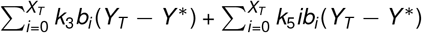 and map

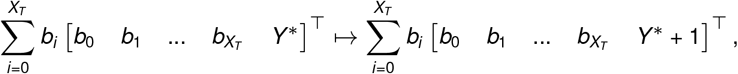

which results in

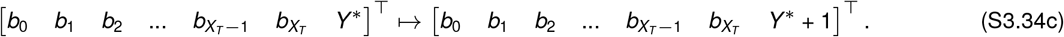
4. State *Y*\* to *Y*\* − 1 occurs with transition intensity 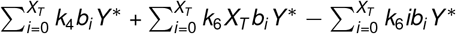 and map

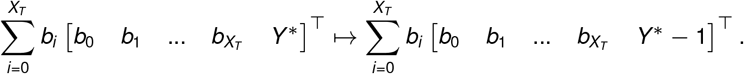

On simplifying, the above map becomes

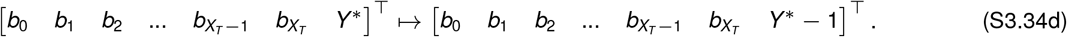

We can now write the dynamics of moments of the form 〈*b_i_Y*\*^*m*^〉 for *m* = 0, 1, 2. Let us begin with 〈*b_i_*〉.

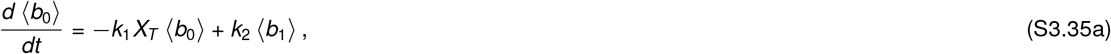

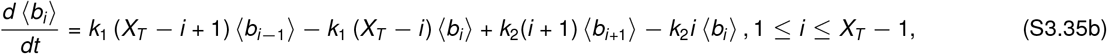

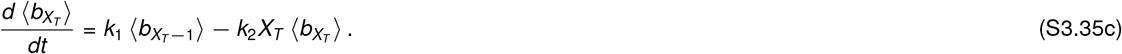

Recalling the definition of *b_i_*, we note that 〈*b_i_*〉 is same as the probability that *X*\* = *i*. We have solved these equations in a slightly different notation in (S3.8). Therefore, the solution to these ODEs is

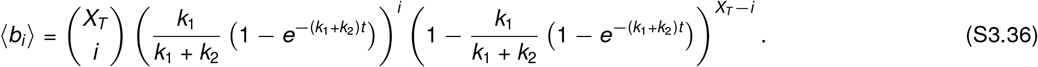

Next, we write the dynamics for 〈*b_i_Y*\*〉.

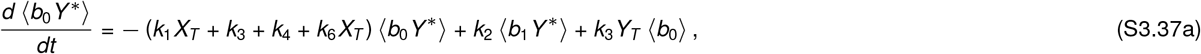

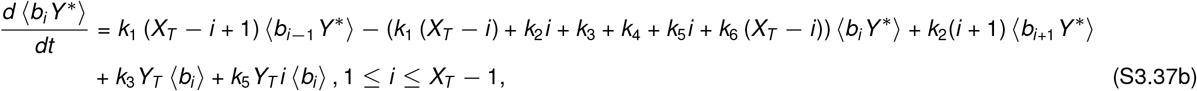

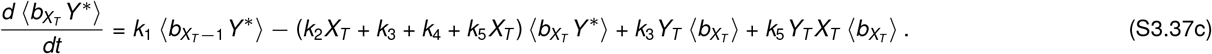

Finally, the ODEs describing the time evolution of 〈*b_i_Y*\*^2^〉 are as follows.

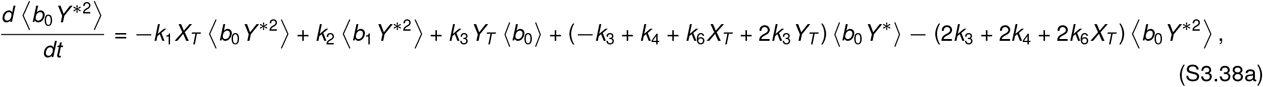

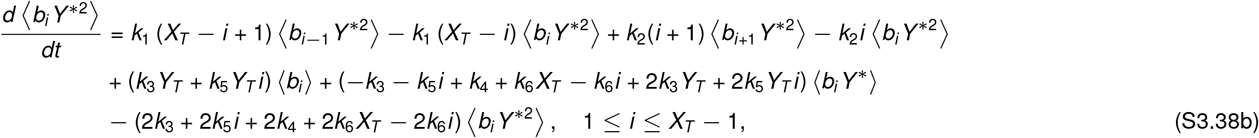

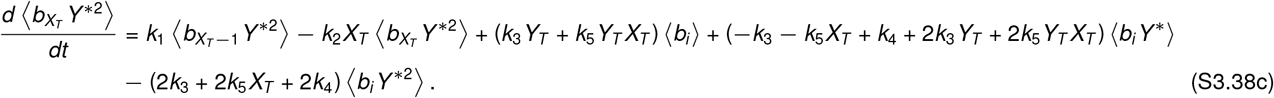

These ODEs require initial condition to compute transient moments which we discuss below.

###### Setting initial condition

In absence of stimulus, we have that 〈*b*_0_〉=1, because no receptors should be active. All other 〈*b_i_*〉 = 0. Furthermore, 〈*b_i_Y*\*〉 = 〈*b_i_*〉 〈*Y*\*〉 and 〈*b_i_Y*\*^2^〉 〈*b_i_*〉 〈*Y*\*^2^〉. Therefore the mean and the second moment at time *t* = 0 are given by the first two moments of the Binomial distribution with parameters 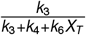 and *Y_T_*. Therefore, the initial condition is

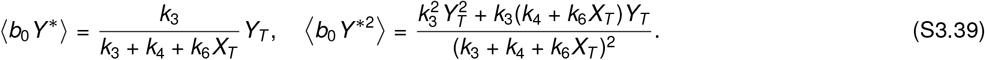

###### Semi-analytical solution using linear algebra

Let 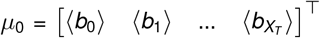 be the collection of the moments of *b_i_*. Then the ODEs can be compactly written as

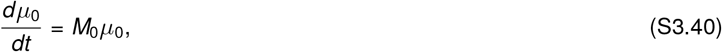

which has the solution 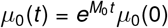. We also note that ∑_*i*_〈*b_i_*〉 = 1 at all times.

The matrix *M*_0_ is tridiagonal, but its inverse does not exist. This does not affect computation of the transient solution as long as we respect the constraint that all 〈*b_i_*〉 sum up to one. For steady-state solution, however, we have to solve

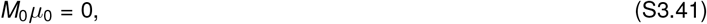

which only exhibits a trivial solution *μ*_0_ = 0. To force the summation requirement, we reduce the system such that we get rid of the last equation corresponding to 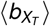. We then substitute 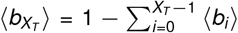 wherever we have 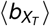. This gives us a reduced system of equation

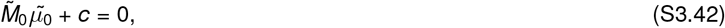

which can be straightforwardly solved using standard linear algebra tools.

It is important to note that we already know the transient as well as the stationary solution for these equations – since 〈*b_i_*〉 are probabilities. However, we present the linear algebra approach for completeness. We will this approach to compute the higher order moments for which analytical solutions are not known.

Let us now solve for the moments 〈*b_i_Y*\*〉. To this end, we collect all the required moments in *μ*_1_ defined as

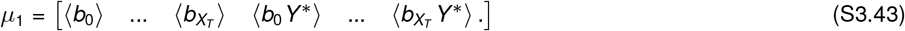

The corresponding ODE system is then

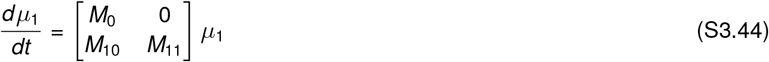

As before, we can now compute the solution using matrix exponential. For the moments 〈*b_i_ Y*\*^2^〉, we can similarly define *μ*_2_

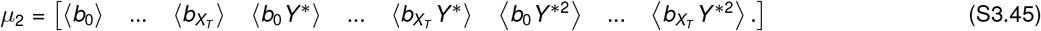

Then we can write the ODE system:

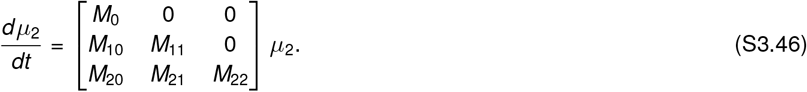

### S4 Effect of receptor internalization

#### S4-A Simple model

Let us begin with a simple model that includes the production of inactive receptors with rate *k_p_*, removal of inactive receptors with rate *k_d_* and removal of active receptors of 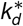. The ODE model for the set up is

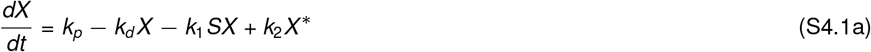

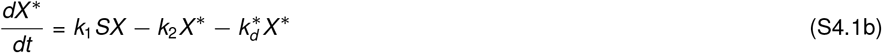

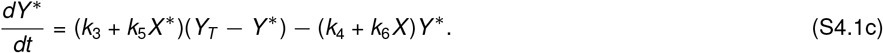

Let us first determine the initial condition before the stimulus arrives. In this case, we have

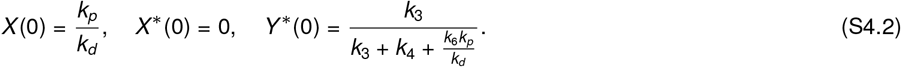

Also, the steady-state solution is

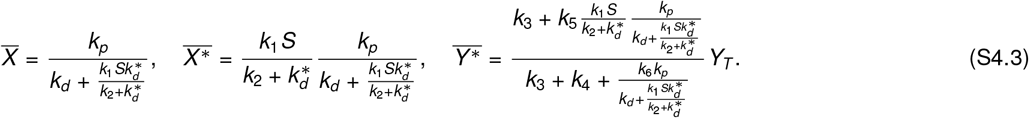

**Figure S4.1:**
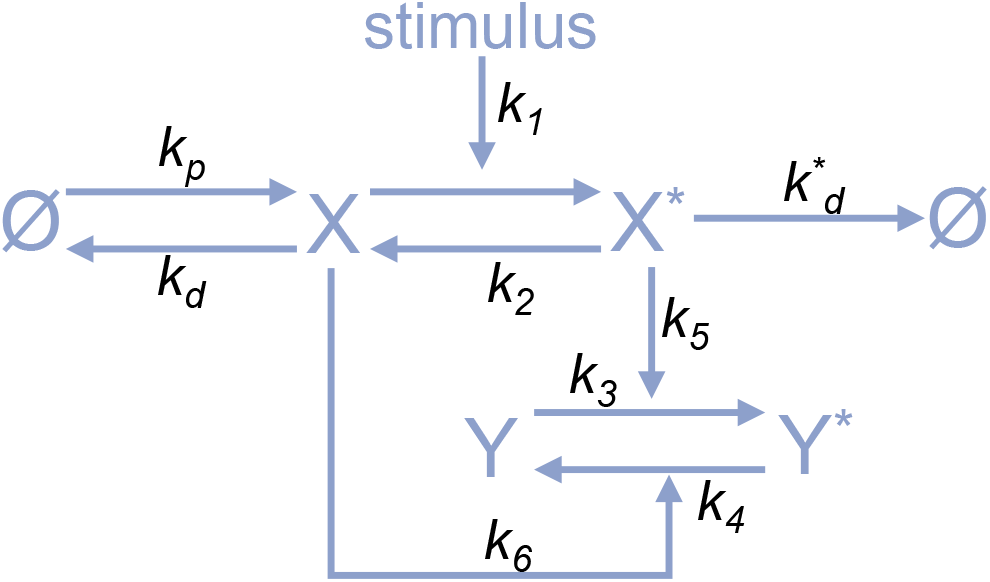
Concerted mechanism with receptor production and degradation.

#### S4-B Solution to receptor dynamics

Our goal here is to examine the effect of receptor removal on different signaling mechanisms. To that end, let us first compute the dynamics at the receptor level.

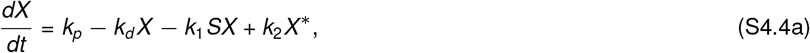

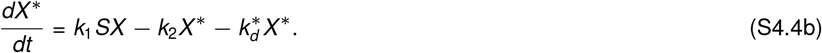

Let 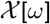 and 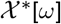 respectively denote the Laplace transforms of *X*(*t*) and *X*\*(*t*). Taking the initial conditions as 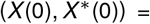 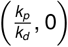 the Laplace transforms of above ODEs results in the following algebraic relations

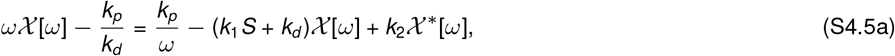

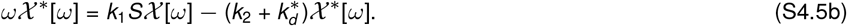

The solution for 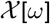 and 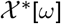 is

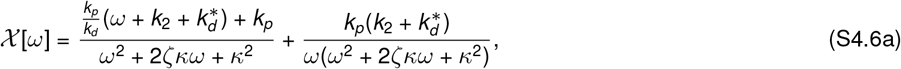

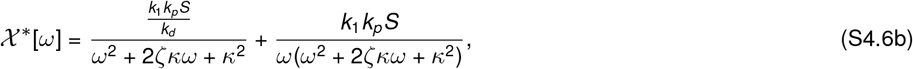

where we have used the following notation

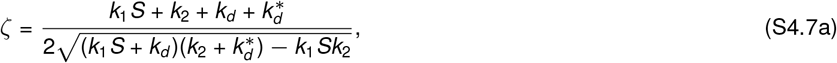

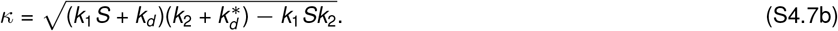

The roots of the term *ω*^2^ + 2*ζκω* + *κ*^2^ are

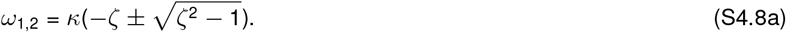

The following usual relations hold for *ω*_1_ and *ω*_2_:

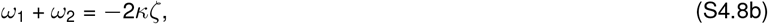

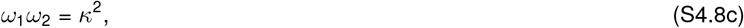

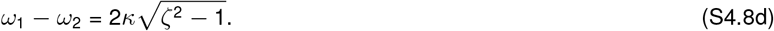

It is easier to take the inverse Laplace transform of 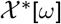 in order to compute *X*\*(*t*):

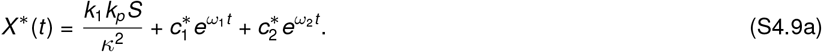

Here the terms 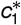 and 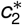 are

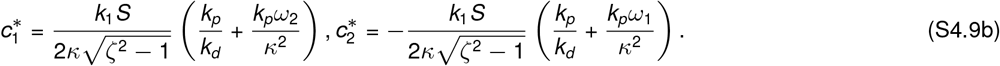

Using the solution of *X*\*(*t*), *X* (*t*) can also be computed as follows.

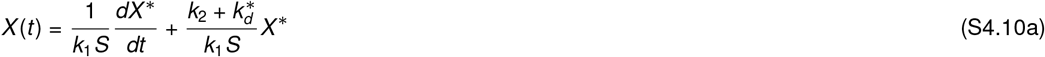

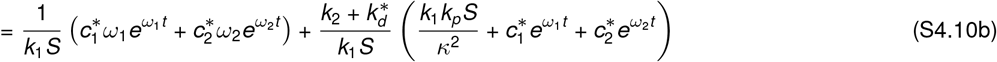

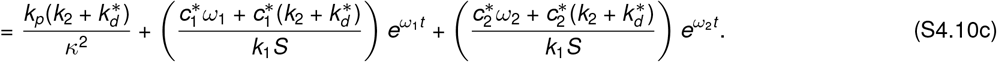

Having determined these solutions, we next provide a lower bound on *ζ*. It is worth noting that *ζ* > 1 implies that the roots *ω*_1,2_ are real.

##### A lower bound for *ζ*

The parameter *ζ* defined in (S4.7) is always greater than one, regardless of the choice of parameters. To see this, we look at *ζ*^2^

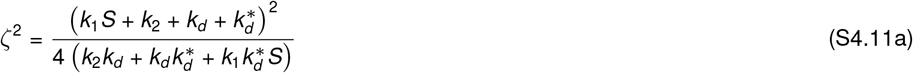

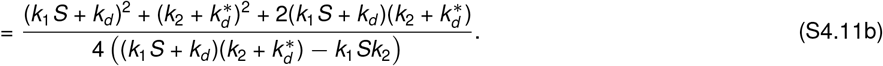

This implies that

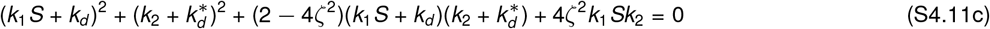

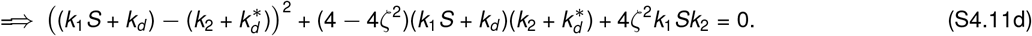

Because all terms in the above equation are positive, except may be for 4 − 4*ζ*^2^, a real solution for *ζ* exists only if 4 − 4*ζ*^2^ < 0. Therefore, *ζ* > 1. Consequently, the roots *ω*_1_ and *ω*_2_ defined in (S4.8a) are negative and satisfy

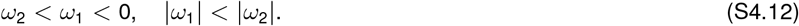

#### S4-C Effect of relative timescales

It is noteworthy that both *X* and *X*\* have two timescales for relaxing to their respective steady-states, determined by *ω*_1_ and *ω*_2_. Because |*ω*_2_| > |*ω*_1_|, we refer to the timescale set by *ω*_2_ as fast timescale and the one set by *ω*_1_ as the slow timescale. The parameter *ζ* controls the difference between the magnitudes of *ω*_1_ and *ω*_2_. What is the impact of these two timescales on the trajectories of *X* (*t*) and *X*\*(*t*)?

Let us first consider *X*\*(*t*) given by (S4.9a). At *t* = 0, the trajectory begins from *X*\*(0) = 0. Consider a scenario where *ζ →* 1, implying that *ω*_1_ ≈ *ω*_2_

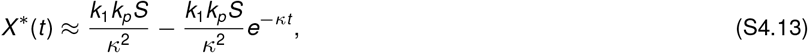

which increases over time to reach the steady-state 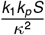. Suppose that *ζ* is now increased. The terms 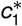 and 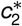 relax with different timescales. Specifically, 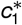 relaxes at a slower timescale than 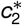. Because 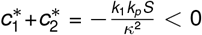, at least one of has to be positive. If we choose a large *ζ* such that at a small time *t′*, the contribution from 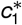 does not change whereas 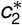 term reaches its “quasi-stationary” value. The solution for *t* < *t′* can then be approximated as

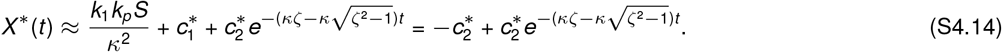

If we assume that 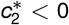, then the quasi-stationary solution is given by

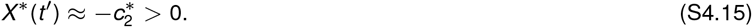

Although this analysis is not rigorous, it equips us with requirements to obtain a response that first attains a peak value above its final steady-state value. Specifically, we need that the coefficient 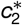 that multiplies the fast timescale exponential term be negative and its magnitude should be greater than the final steady-state. In other words, we need:

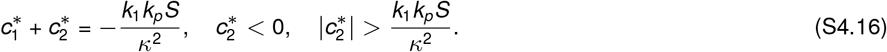

We substituted the values of *ζ* and *κ* from (S4.7) and used to symbolic solver to solve the above inequalities. We obtain that the following should be satisfied:

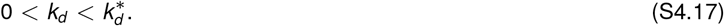

We get similar requirements for a trajectory of *X* (*t*) that starts from *X* (0) = *k_p_/k_d_*, then decreases with a fast timescale below its final stead-state value (i.e., attains a quasi-stationary value) and then relaxes back to the steady-state value. These conditions are

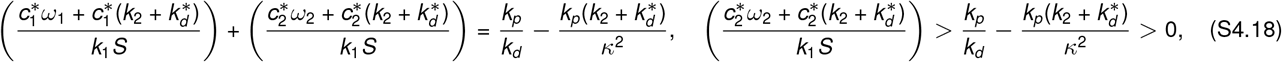

As before, substituting the expressions of *ζ* and *κ* shows that these requirements are same as having 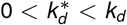.

### S5 Alternating activation and derepression

In this section, we consider signaling cascades consisting of alternating activation and derepression based switches. The first cascade is shown in Fig. S5.1(a). It is built upon the activation mechanism of Fig. 1(a) in the main text, where the receptor activates a downstream switch (*Y* ⇄ *Y*\*). We add a downstream switch (*Z* ⇄ *Z*\*) which is derepressed. The second cascade, shown in Fig. S5.1(b), is a modification of the derepression mechanism of Fig. 1(b) in the sense that a downstream component is now activated by the derepressed switch.

#### S5-A Activation followed by derepression

The ODEs that govern the dynamics of this cascade are

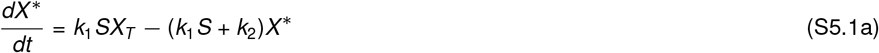

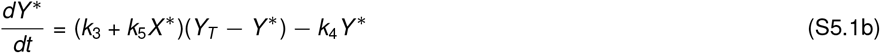

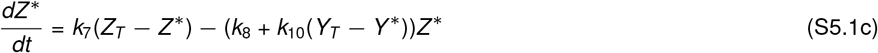

We obtain the steady-states by setting each of the derivatives to zero. We express each of the steady-states in a similar form as that of (3) in the main text

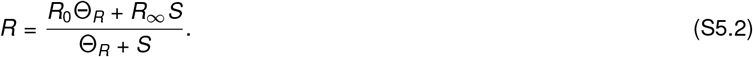

For example, steady-state of *X*\* is specified by

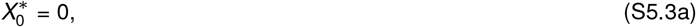

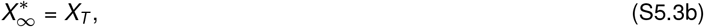

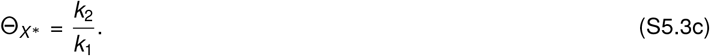

**Figure S5.1:**
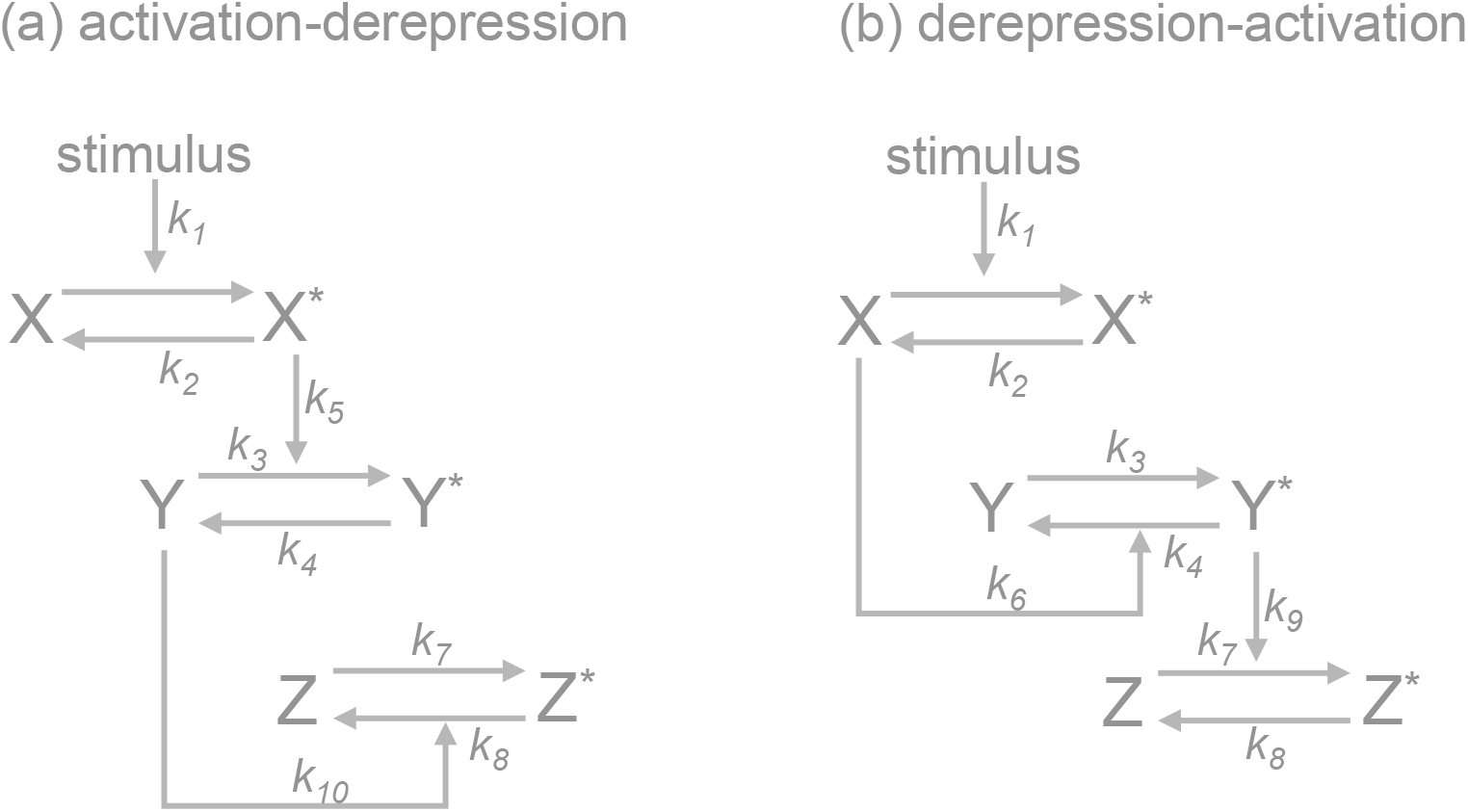
Three tier cascades with alternating activation and derepression mechanisms

The steady-state of *Y*\* is specified by

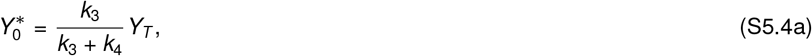

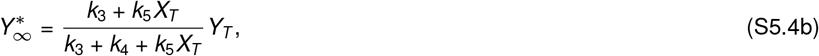

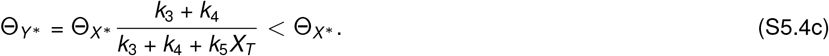

As expected, activation caused the dose-response of 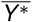 to shift towards left in comparison with that of 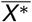, i.e., Θ_*Y*\*_ < Θ_*X*\*_. Finally, the steady-state of *Z*\* is specified by

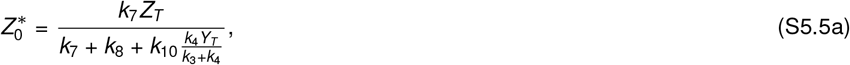

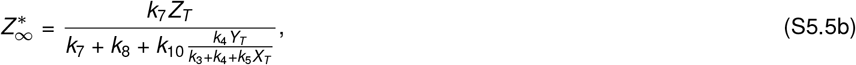

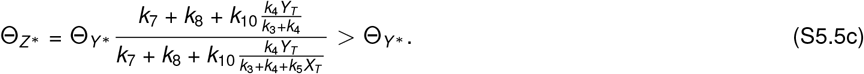

We observe that Θ_*Z*_ > Θ_*Y*\*_. This means that the derepression layer has an opposite effect of activation and shifts the dose-response back towards right.

#### S5-B Derepression followed by activation

The ODEs that govern the dynamics of this cascade are

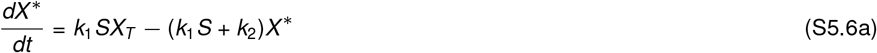

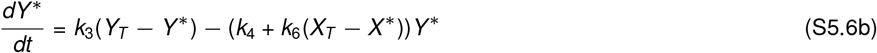

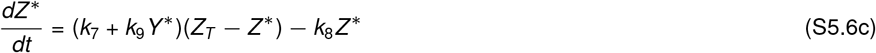

For this model, the steady-state of 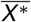 has the same specification as (S5.3). The steady-state 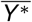 is prescribed by

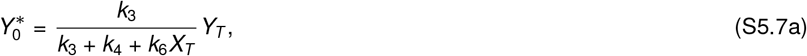

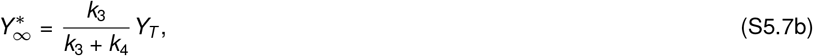

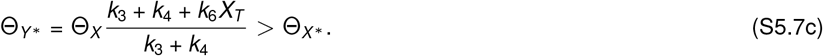

Because Θ_*Y*\*_ > Θ_*X*\*_, the dose response of *Y*\* is towards the right to that of *X*\*. This results from the fact that this switch is governed by a derepression mechanism. We now look at the parameters specifying 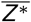:

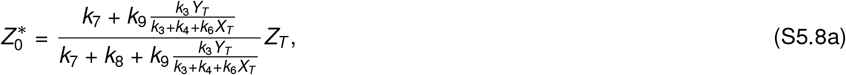

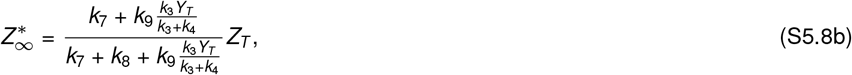

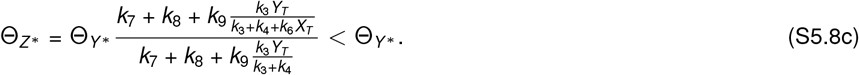

We see that Θ_*Z*\*_ < Θ_*Y*\*_. So, the dose-response of *Z*\* is towards the left of *Y*\*, which implies that activation of the third layer counteracts the shifting caused of derepression of the second layer. It is important to point out that the effects of these mechanisms on 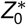 and 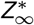 are different. A systematic analysis of these effects on alternating cascades will be carried out in a future work.

